# Divergent transcriptional architectures beyond core CAM genes in facultative and constitutive CAM species in Tillandsia

**DOI:** 10.1101/2024.05.09.593278

**Authors:** Clara Groot-Crego, Sarah Saadain, Marylaure de La Harpe, Jaqueline Hess, Michael H.J. Barfuss, Walter Till, Gert Bachmann, Wolfram Weckwerth, Christian Lexer, Ovidiu Paun

## Abstract

Crassulacean acid metabolism (CAM) is a water-efficient photosynthetic strategy involving a coordinated suite of complex traits including metabolic, anatomical and regulatory aspects that shift across the diel cycle. While CAM has evolved repeatedly in land plants, the evolutionary routes enabling this convergence remain elusive. Whereas the same core CAM (de)carboxylation genes are consistently involved, a key question is whether distinct CAM phenotypes also depend on a shared set of auxiliary genes, reflecting a quantitative continuum of expression, or whether they can instead emerge through divergent or redundant peripheral solutions. The bromeliad subgenus *Tillandsia*, with diverse photosynthetic strategies, offers an ideal system to explore this question. Using physiological and transcriptomic analyses of well-watered and water-limited accessions of two closely related species, we characterized facultative and constitutive CAM. By comparing orthologous gene expression and orthogroup recruitment, we found that while both species performed CAM upon water-withholding, transcriptional shifts in pathways related to stomatal movement, sugar/malate transport, aquaporins and starch metabolism showed minimal overlap. Core enzymes involved in the CAM (de)carboxylation cycle exhibited broadly shared expression patterns, yet the facultative CAM species uniquely up-regulated PPC2 at night instead of the canonical CAM-related PEPC ortholog PPC1. Our study reveals that, while the expression of certain core CAM enzymes is conserved, the surrounding transcriptional architecture can differ substantially even between closely related species. This supports a model in which CAM evolves through a mosaic recruitment of functionally equivalent, yet non-orthologous genes - underscoring its flexible and modular genetic architecture. These insights advance our understanding of the mechanisms enabling the repeated evolution of CAM and its capacity to facilitate adaptive diversification.

## 2. INTRODUCTION

The repeated evolution of complex traits has been a long-standing matter of study for evolutionary biologists. It remains unclear how traits which need coordinated changes across multiple organismal layers can evolve frequently and repeatedly across the tree of life. One explanation is that similar phenotypes can arise through different sets of mutations, also referred to as “genetic routes”, assuming complex traits are polygenic and the underlying genes show substantial redundancy (Barghi et al., 2020; Goldstein & Holsinger, 1992; Láruson et al., 2020; M. A. Nowak et al., 1997). Indeed, convergent phenotypes lacking a shared genetic basis have been documented across a broad spectrum of life including beach mice (Steiner *et al*., 2009), conifers (Yeaman *et al*., 2016), lab-reared *Drosophila* (Barghi *et al*., 2019), cacao plants (Hämälä *et al*., 2020) and *Heliosperma* ecotypes (Szukala et al., 2023).

Crassulacean acid metabolism (CAM) represents a prime example of a collection of coordinated, complex yet highly evolvable traits, having evolved independently at least 66 times across 38 plant families (Gilman *et al*., 2023). CAM plants reconfigure photosynthetic metabolism across the diel cycle by assimilating CO_2_ at night rather than during the day, as C_3_ plants do. Nocturnally-fixed CO_2_ is then temporarily converted to malate by phosphoenolpyruvate carboxylase (PEPC) and malate dehydrogenase (MDH). Malate is stored in vacuoles and later decarboxylated during the day to supply CO_2_ for the Calvin-Benson-Bassham cycle. This enables daytime stomatal closure, increasing Rubisco efficiency by raising intracellular CO_2_ levels (Osmond, 1978) and drastically reducing water loss through evapotranspiration (Borland *et al*., 2014). These adaptations are especially advantageous under high temperatures or drought stress, which exacerbate Rubisco oxygenation (Heyduk *et al*., 2021).

CAM integrates several interdependent traits, including temporally coordinated transcriptional regulation to set the alternate biochemical pathway in motion, and cellular processes like stomatal movement, sugar and malate transport, and starch metabolism (Borland et al., 2016; Schiller & Bräutigam, 2021). Additionally, certain anatomical features often appear associated with CAM, leading to the hypothesis that they may be a prerequisite for a plant to fully express CAM (Nelson *et al*., 2005). Examples of such anatomical features are enlarged vacuoles (Heyduk et al., 2016; Males, 2018), tighter packed mesophyll cells (Nelson & Sage, 2008) and succulence (Gibson, 1982; Griffiths *et al*., 2008), though for the latter, intra-specific variation does not appear correlated with CAM (Martin *et al*., 2009).

CAM was initially described as having a discrete, bi-modal distribution of phenotypes, yet detailed physiological studies of C_3_ species with CAM relatives have revealed weak and facultative forms of CAM that occur without major anatomical changes (Heyduk et al., 2021; Neales, 1975; Pierce et al., 2002; Silvera et al., 2005; Winter & Holtum, 2002). These findings prompted a shift towards conceptualizing CAM as a continuum, though debate persists (Bräutigam et al., 2017; Schiller & Bräutigam, 2021; Winter & Smith, 2022). The CAM continuum concept remains difficult to reconcile with the many discrete CAM categories in the literature (e.g. facultative, weak, strong, constitutive CAM, CAM-cycling (Winter, 2019)), which do not clearly align with a continuous phenotype spectrum (Edwards, 2023). Additionally, weak/facultative CAM plants can be difficult to detect, as carbon isotope ratios, the most cost-effective method to measure CAM activity on a large taxonomic scale, lack sensitivity for these phenotypes (Pierce et al., 2002; Winter & Holtum, 2002). More detailed measurements, such as titratable acidity or gas exchange under standard and drought conditions, are needed to distinguish weak/facultative CAM from C_3_ species.

The evolutionary trajectory of CAM and the molecular similarity across CAM phenotypes remain unclear. While diel expression shifts of core CAM enzymes such as PEPC appear broadly conserved among CAM plant taxa, the realized CAM phenotype likely involves numerous small-effect genes stemming from broader metabolic, regulatory and anatomical networks, potentially with a high redundancy. Comparative analyses between different CAM phenotypes can reveal whether CAM evolution follows a constrained, shared evolutionary path or instead involves large genetic redundancy. For example, Heyduk *et al*. (2018) found both shared and variable expression patterns in CAM-related genes among weak and strong CAM phenotypes in Agavoideae, but more studies comparing the molecular basis of CAM between closely related taxa are needed to obtain a comprehensive view of genetic redundancy and constraints in CAM evolution.

The subgenus *Tillandsia* (genus *Tillandsia*, Bromeliaceae) is an emerging model for studying repeated and rapid evolution of key innovation traits. As part of one of the fastest diversifying plant lineages (Givnish *et al*., 2014), the subgenus comprises an adaptive radiation of over 250 species formed within the last 7 million years (Barfuss et al., 2016; Mendoza et al., 2017; Yardeni et al., 2025). Numerous key innovations characterise this radiation, including CAM, which spans a broad phenotypic range from C_3_-like to strong constitutive CAM (Crayn *et al*., 2004; De La Harpe *et al*., 2020).

In this study, we compare a closely related species pair exhibiting facultative and constitutive CAM to assess the degree of genetic redundancy underlying this suite of interdependent complex traits. We consider CAM as determined by many interacting genes, cellular processes, and pathways. If CAM evolution is tightly constrained, both phenotypes should represent positions along a shared evolutionary trajectory, mediated by the same set of genes but expressed to different degrees. Conversely, limited overlap in gene expression, at least in some parts of the network, would suggest a (partly) redundant genetic architecture, consistent with greater evolvability and multiple independent evolutionary routes to CAM evolution.

The species *T. vanhyningii* Harms (formerly *T. ionantha* var. *vanhyningii* L.B.Sm., elevated to species level by Beutelspacher & Garcia-Martinez (2021)), and *T. leiboldiana* Schltdl. are closely related members of the subgenus *Tillandsia,* with a maximum divergence time of 3.2 Mya (Barfuss et al., 2016; Mendoza et al., 2017; Yardeni et al., 2025). *Tillandsia vanhyningii* is a succulent “grey” species with dense trichome cover, whereas the “green” *T. leiboldiana* typically forms an impounding tank and is comparatively less succulent. These species display some of the most disparate carbon isotope values in the subgenus (De La Harpe *et al*., 2020), with *T. vanhyningii* having δ^13^C values typical of strong, constitutive CAM (-13.9), while *T. leiboldiana* shows values typical of C_3_ photosynthesis (-31.3). However, recent metabolomic, genomic, and transcriptomic analyses suggest that *T. leiboldiana* may be a facultative CAM species (Groot Crego *et al*., 2024). Despite minimal night-time malate accumulation, *T. leiboldiana* shows CAM-like expression in certain CAM-related genes, including PEPC kinase, under well-watered conditions, indicating a latent CAM capacity potentially activated under drought-induced stress.

We subjected accessions of both species to standard and water-limited conditions, and measured titratable acidity, gas exchange and transcriptomic profiles across the diel cycle to compare their photosynthetic phenotypes under both watering regimes (Fig. 1). Our study characterizes their photosynthetic phenotypes and elucidates how water-withholding modulates CAM expression. We show that, although both species perform CAM under water-limited conditions, they exhibit substantial divergence in transcriptional modulation of CAM-related, peripheral genes beyond the core CAM genes. These findings illuminate the degree of genetic redundancy underlying CAM and provide insight into the multiple evolutionary routes by which CAM can arise.

**Figure 1.**
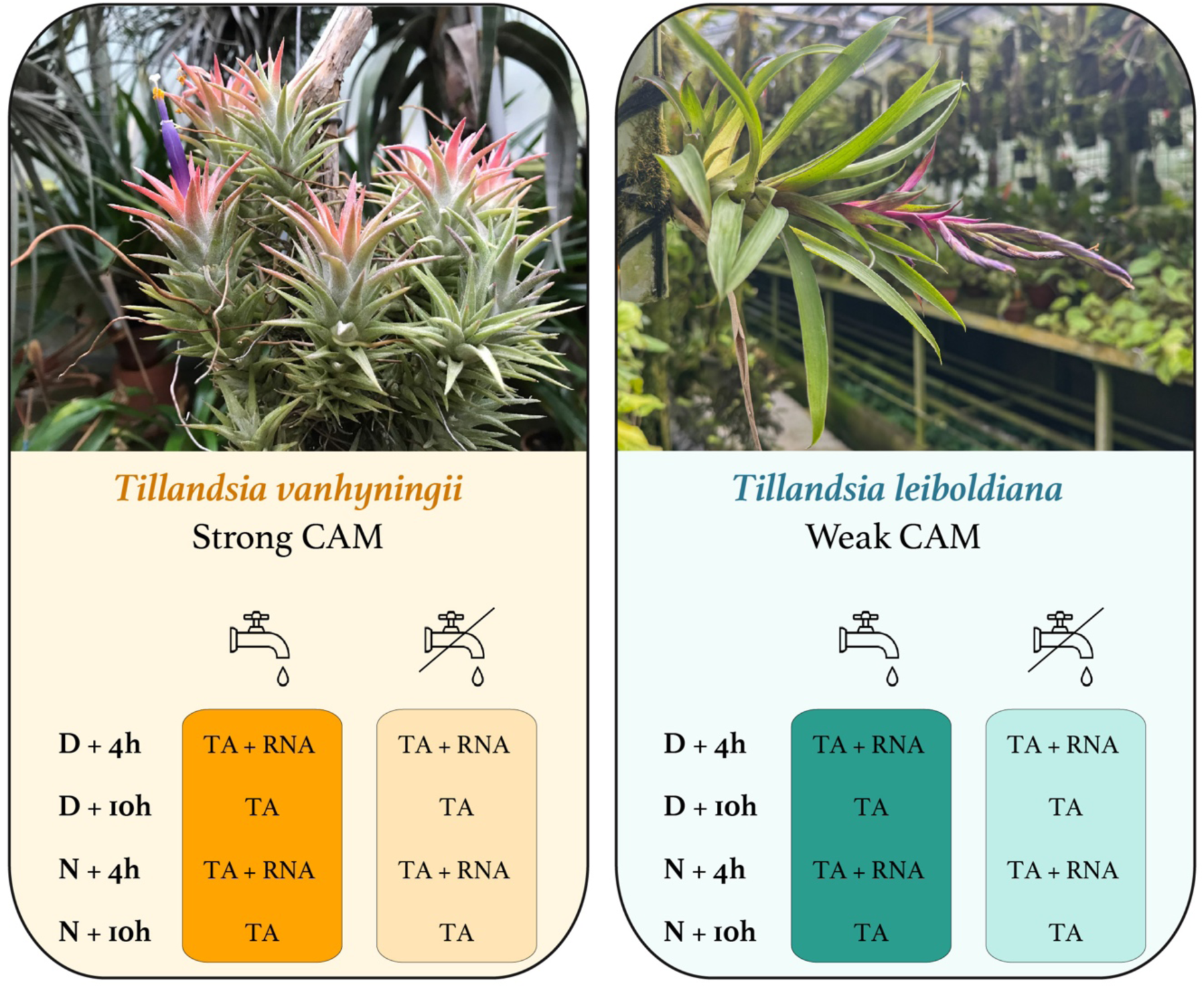
Overview of species and experimental set-up. Left, Tillandsia vanhyningii (Photograph by Michael H.J. Barfuss, 2023); right, Tillandsia leiboldiana (Photograph by Manuel Lasserus, 2024), both at the Botanical Garden of the University of Vienna. Experimental setup, consisting of sampling at four time points that are six hours apart: 4 and 10 hours after dawn (D), plus 4 and 10 hours after nightfall (N). Titratable acidity (TA) was quantified for each condition, whereas RNA-seq data was analysed for the time points indicated with RNA. A separate, additional drought experiment was performed on T. leiboldiana (not shown here) to study net CO_2_ uptake across diel cycles, to further phenotype its photosynthetic metabolism.

## 3. MATERIALS AND METHODS

### 3.1. Experimental set-up and sampling

We designed an experiment to test the respective photosynthetic phenotypes of *T. vanhyningii* and *T. leiboldiana* in standard (SW) and water-limited (WL) conditions. The study comprised six clonal accessions per species (Table S1), which were off-shoots of similar size, as a proxy for age similarity, and were large enough to be considered adult plants able to flower. Accessions from the plant collection of the Botanical Garden of the University of Vienna were placed under identical greenhouse conditions in October 2018. The accessions were allowed to acclimate to the specific greenhouse conditions for six weeks, before being placed in a regime of 12 hours of light / 12 hours of dark for 14 days, by complementing daylight with artificial lights between 6 am and 6 pm. The temperature was maintained at 20 °C during nighttime and 22 °C during daytime. During these 14 days, three accessions of each species were placed in a water-limited regime by complete water withholding. The remaining three accessions were kept in standard watering conditions used to maintain the plants in the botanical garden (1x every two days). After 14 days, all accessions were sampled every 6 hours (Four time points: Dawn + 4 hours (10 am), Dawn + 10 hours (4 pm), Nightfall + 4 hours (10 pm), Nightfall +10 hours (4 am)), resulting in three replicates for the combination of treatments (species x time point x watering regime) (Fig. 1). As every accession was sampled four times sequentially, leaf material was collected at each time point by delicately pulling out an entire leaf from the base of the plant to minimise wounding. The tip and base of the leaf were then removed, and the middle part was immediately placed in liquid nitrogen and stored at -80 °C.

### 3.2. Titratable acidity measurements

Titratable acidity (TA) was measured following Wai *et al*. (2019). Frozen leaf material for all four time points, two watering regimes and two species (48 samples in total) was weighed and ground in a pre-cooled mortar to a powder, then added to a 5 ml solution of 80 % EtOH. Next, the samples were incubated for one hour in an 80 °C water bath with occasional vortexing. Tubes were centrifuged at 5,000 rpm for 5 minutes, after which the supernatant was transferred and the pellet discarded. The extractions were then let to cool to room temperature. Using a pH526 WTM Sentix mic metre, each sample was measured by titrating 0.1 N NaOH to the solution until reaching a pH of 8.2 exactly and recording the volume of added NaOH. TA in μ-equivalents per gram of fresh mass (g FM) was calculated using the formula: TA (µ-equivalents/g FM) = [V_NaOH_ (µL) × C_NaOH_ (µmol/µL)] / m_sample_ (g), where C_NaOH_ is equal to 0.1. Since NaOH is a monovalent base, the number of µ-equivalents of titratable acid (from both mono- and multivalent acids collectively) is equal to the number of micromoles of NaOH consumed during titration.

To test whether TA significantly fluctuated across time within each species and watering regime, we constructed a Linear Mixed-Effects Model (LMM) for each species as follows:

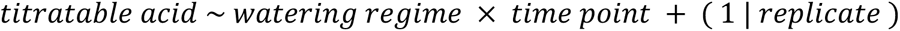

Normality of TA measurements within each species was tested beforehand. In the LMM model, titratable acid represented the dependent variable, while watering regime and time point were fixed effects, and their interaction was included into the model. To control for random variation in TA measurements across replicates of the same time point and watering regime, the model included a random intercept for each replicate. The analysis was performed using the *lmer* function from R package *lme4* (Bates *et al*., 2015). The goodness of fit of the model was evaluated by plotting a QQ-Plot and the fitted values to residuals for each species. The significance of fixed effects was evaluated by obtaining a p-value using the R package *lmerTest* (Kuznetsova *et al*., 2017). Afterwards, a post-hoc Tukey HSD test was performed in the R package *emmeans* to assess the effects of individual watering regimes and time points on TA.

### 3.3. Gas exchange measurements of *T. leiboldiana*

A separate experiment was performed on *T. leiboldiana* in October 2025 to assess net CO_2_ uptake progressively throughout a period of water-withholding. Leaves of one *T. leiboldiana* clone were placed in a gas exchange experiment chamber over a period of eight days where the plant was not watered. The net CO_2_ uptake and other metrics were measured by a WALZ GFS-3000 system. The ambient temperature during the experiment was maintained between 22 and 24 °C, and the relative humidity at 35 % outside the leaf clamp and at 50% inside. Full spectrum LED lights (maximum 340 PPFD) were turned on gradually between 6 and 7 am, and 6 and 7 pm, resulting in a 12-hour light cycle. At the time of finishing the experiment (9 days without watering), the plant showed clear signs of drought-stress (yellowing and curling leaves). The plant was then placed under a 14-day recovery period beginning with full submersion in demineralized water, followed by daily spraying in a greenhouse with same climatic conditions as in the experimental chamber, except for a rH of 70 %. Net CO_2_ uptake was measured again at the end of the recovery period. Due to the small size and thick, concave shape of *T. vanhyningii* leaves, it was not possible to measure gas exchange reliably on this species with this system.

### 3.4. RNA extraction and sequencing

RNA-seq data was obtained for samples at time points 10 am (D+4) and 10 pm (N+4) for two watering regimes and two species (24 samples in total). Total RNA was extracted using the QIAGEN RNeasy® Mini Kit. We evaluated the purity and concentration of the extractions using Nanodrop; RIN and fragmentation profiles were obtained with a BioAnalyzer™ (Table S1). RNA library preparation and poly-A capture were performed at the Vienna Biocenter Core Facilities (VBCF) using a NEBNext stranded mRNA kit. The libraries were sequenced on Illumina NovaSeq SP 150 PE.

### 3.5. RNA-seq data processing

Raw sequence quality was assessed using FastQC v.0.11.8 (https://www.bioinformatics.babraham.ac.uk/projects/fastqc/) and MultiQC (Ewels *et al*., 2016). Adapters and low-quality bases were trimmed off using AdapterRemoval v.2.3.1 (Schubert *et al*., 2016) with settings --trimns --trimqualities --minquality 20 --trimwindows 12 --minlength 36. The trimmed *T. leiboldiana* data was aligned to its conspecific reference genome (GenBank accession: GCA_029204045.2, Groot Crego et al 2024), while the *T. vanhyningii* data was aligned to the reference genome of its closest relative, *T. fasciculata* (GenBank accession: GCA_029168755.2, Groot Crego et al. (2024)). Plastid phylogenies show that *T. fasciculata* and the *T. ionantha* complex, which includes *T. vanhyningii* (both constitutive, strong CAM), diverged no more than 1.8 mya (Vera-Paz *et al*., 2023). Yardeni et al. (2025) reported 0.008 substitutions per site between the two lineages based on a maximum-likelihood inference using whole-genome data. Given this close evolutionary proximity, we consider these species suitable for cross-species mRNA read alignment. These were performed with STAR v. 2.7.3a (Dobin *et al*., 2013) in one pass using a GFF file to specify exonic regions. To demonstrate that using a non-conspecific reference genome does not substantially influence our findings, we provide in supplementary a comparison of analyses on *T. leiboldiana* mapped to its own reference genome, versus to *T. fasciculata*.

### 3.6. Differential gene expression analysis

Reads were quantified for paired-end and reversely stranded reads (options -p, -s 2) in the subread package FeatureCounts v.2.0.6 (Liao *et al*., 2014). Counts were generated by counting reads per exon, yet aggregating counts across exons for each gene using FeatureCounts’ in-built meta-feature function. Differential gene expression (DGE) analyses were performed for each species separately using the Bioconductor package edgeR v.4.2.1 (Robinson *et al*., 2009).

We built a design matrix incorporating the two investigated treatments (time point and watering regime). Genes where three or fewer samples of the same species had a Counts Per Million (cpm) value smaller or equal to 1 were filtered out. Library size normalisation was done using edgeR’s default trimmed mean of M-values (TMM) method (Robinson & Oshlack, 2010). The Cox-Reid profile-adjusted likelihood (CR) method was used to estimate dispersion. Genes were tested for differential expression between time points and between watering regimes separately for each species by fitting a negative binomial General Linear Model (GLM) with the following contrasts: Day versus Night at SW = Day_SW - Night_SW, Day versus Night at WL = Day_WL - Night_WL, WL vs SW at day = WL_day - SW_day, WL vs SW at night = WL_night - SW_night.

To test the fit of these GLMs, we conducted a quasi-likelihood F-test. The level of significance was adjusted with the Benjamini-Hochberg False Discovery Rate (FDR) correction to account for multiple testing. Genes were considered differentially expressed (DE) when the FDR-adjusted p-value was below 0.05 and the expression had an absolute log-fold change of at least 1.5. The number of DE genes for each treatment (time point and watering regime) was then visualised and compared between species using ggplot2.

### 3.7. Overlap in transcriptomic response

To understand the extent of overlap in the transcriptomic response to time and watering regime between species, DE genes of both species were studied on the orthogroup level, using orthology information of *T. leiboldiana* and *T. fasciculata* from Groot Crego et al. (2024). Orthogroups containing a DE gene in at least one species are hereafter referred to as DE orthogroups. The percentage of DE orthogroups containing more than one DE gene in a single species were calculated for each comparison to understand the relationship between gene-level and orthogroup-level differential expression (reported in Table S8). Venn diagrams were computed to visualize overlaps in DE orthogroups between species using the R package VennDiagram (Chen & Boutros, 2011).

We also assessed overlap in DE orthogroups across time- and watering-dependent comparisons, as CAM-related genes are likely DE in both contexts, particularly in facultative CAM species like *T. leiboldiana*. The overlap in DE orthogroups between comparisons was visualized with UpSetR (Conway *et al*., 2017).

In the discussion of results regarding the transcriptomic overlap between species, timepoints and watering regimes, a distinction is made between within-species comparisons, where the overlap is discussed on the gene level, and between-species comparisons, where it is discussed on the orthogroup level.

### 3.8. Functional interpretation of DE genes

Gene Ontology (GO) enrichment analyses were performed for both sets of DE genes (between time points and watering regimes) within both species using the Bioconductor package topGO v.2.50.0. (Alexa *et al*., 2006). Significance of enrichment was evaluated with the Fisher exact test and the algorithm “weight01”. The top 100 significant GO terms were then compiled sorted by decreasing significance using a custom-made script available at the repository https://github.com/cgrootcrego/Tillandsia_CAM-Drought.

GO enrichment analyses were also performed on each subset of orthogroups representing a section in the Venn-diagrams with at least 25 orthogroups. GO enrichments were performed on the DE genes belonging to the orthogroups listed in each Venn-diagram intersection. For species-specific intersections, annotations from the respective species were used (e.g., *T. leiboldiana* gene annotations for its unique intersections). For overlapping intersections, *T. fasciculata* annotations were used, assuming overlapping annotations between species within orthogroups.

Complementary to a GO enrichment analysis, we identified genes previously described as related to CAM, starch metabolism and glycolysis following Groot Crego et al. 2024. Briefly, *A. comosus* gene IDs related to CAM, starch metabolism or glycolysis compiled by Yardeni et al. (2021) based on a diverse set of studies was used to obtain the corresponding *T. fasciculata* and *T. leiboldiana* orthologs using orthology information from Groot Crego et al. 2024. This resulted in a total of 818 and 753 genes respectively, across 607 orthogroups (Table S2). The orthogroups from this subset that contained DE genes either between time points or watering regimes were then listed by Venn-diagram intersection in Tables S8 and S9.

The abundance of RNA-seq reads for a select group of 53 CAM-related orthogroups was plotted as a heatmap using the R package ComplexHeatmap (Gu *et al*., 2016) for all 24 samples. This subset included 7 orthogroups containing the pineapple genes putatively described as part of the CAM carbon-fixation module by Ming et al. (2015), regardless of their differential expression status. The remaining 46 orthogroups comprised aquaporins (5), circadian clock regulators (11), malate transporters (6), photosystem-related genes (5), vacuolar proton pumps (9), genes related to stomatal movement (4), sugar transporters (5) and one vacuolar invertase (Table S3). Reads were normalised with the TPM (Transcripts Per Million) method using total exonic length of each gene, and converted to a log10 scale, then mean-centred with a z-score transformation. For homologs of the *PEPC* and *PPCK* genes, and other core CAM pathway genes, expression in TPM was additionally visualized by line plots.

A separate subset of orthogroups was selected to investigate the drought-related response to water-limited conditions in both species, by including all DE genes that were annotated with GO term “GO0009414; water deprivation” or belonged to the drought resistance category as specified by Yardeni et al. (2021). This resulted in 32 orthogroups underlying ABA biosynthesis (1), ABA signalling (4), anti-oxidation and ROS (Reactive Oxygen Species) scavenging (5), drought tolerance (7), Late Abundance Embryogenesis proteins (8), inositol biosynthesis (3), jasmonic acid (JA) biosynthesis (1), molybdate ion transport (1) and general stress response (2) (Table S4). A heatmap of their expression values in TPM was constructed as detailed above.

### 3.9. Gene trees of PPC1, PPC2 and PPCK

We identified homologs of the two subfamilies of PEPC, PPC1, PPC2 and of PPCK by running BLASTP on the full set of gene models of *A. comosus* (V3), *Sorghum bicolor* (v.3.1.1), *Oryza sativa* (v.7.0), *Zea mays* (RefGen v.4), *Asparagus officinalis* (v.1.1) and *Arabidopsis thaliana* (TAIR10) available on Phytozome (https://phytozome-next.jgi.doe.gov/), with the PEPC and PPCK gene models of *T. fasciculata* and *T. leiboldiana* as query (Groot Crego *et al*., 2024). Hits were filtered based on identity and target length for each respective gene. For PPC1, we selected all targets longer than 500 aa and with more than 80% identity, resulting in 56 protein sequences across the seven species. For PPC2, the identity threshold was lowered to 70 %, resulting in eight sequences. For PPCK, we selected all sequences with an identity greater than 60 % regardless of target length, resulting in 17 sequences.

Protein sequences were aligned using mafft v.7.520 (Katoh *et al*., 2002) with options *–op 1.5 --ep 0.1 --maxiterate 1000 --localpair*. Gene trees were constructed using IQTREE v.2.2.5 (Minh *et al*., 2020) with options *-B 1000 -T AUTO*. The resulting consensus tree was used for visualisation, collapsing branches leading to Poaceae genes. Nomenclature of PEPC homologs follows Deng et al. (2016) and Heyduk et al. (2022).

## 4. RESULTS

### 4.1. Titratable acidity measurements

In standard conditions (SW), TA displayed a CAM-like diel curve in *T. vanhyningii*, with overnight acid accumulation followed by daytime depletion (Fig. 2a, Table S5). Our LMM model showed that TA significantly fluctuated between time points both under standard and water-limited (WL) conditions, yet there was no significant difference in TA values between watering regimes at the same time point (Fig. 2, Fig. S1, Table S6). In *T. leiboldiana*, TA did not significantly accumulate overnight under SW (Fig. 2b). Under WL, TA values between D+10 and N+10 were significantly different, indicating a slight fluctuation in leaf acidity (Fig. 2b). *T. leiboldiana* exhibited a markedly different temporal curve in TA accumulation between SW and WL, as TA values between watering regimes at each time point was significantly different in all but one time point (Fig. 2a, Table S6). This indicates a weak, facultative CAM cycle in *T. leiboldiana*, whereas *T. vanhyningii* exhibited temporal TA signatures that are characteristic of a strong, constitutive CAM plant.

**Figure 2:**
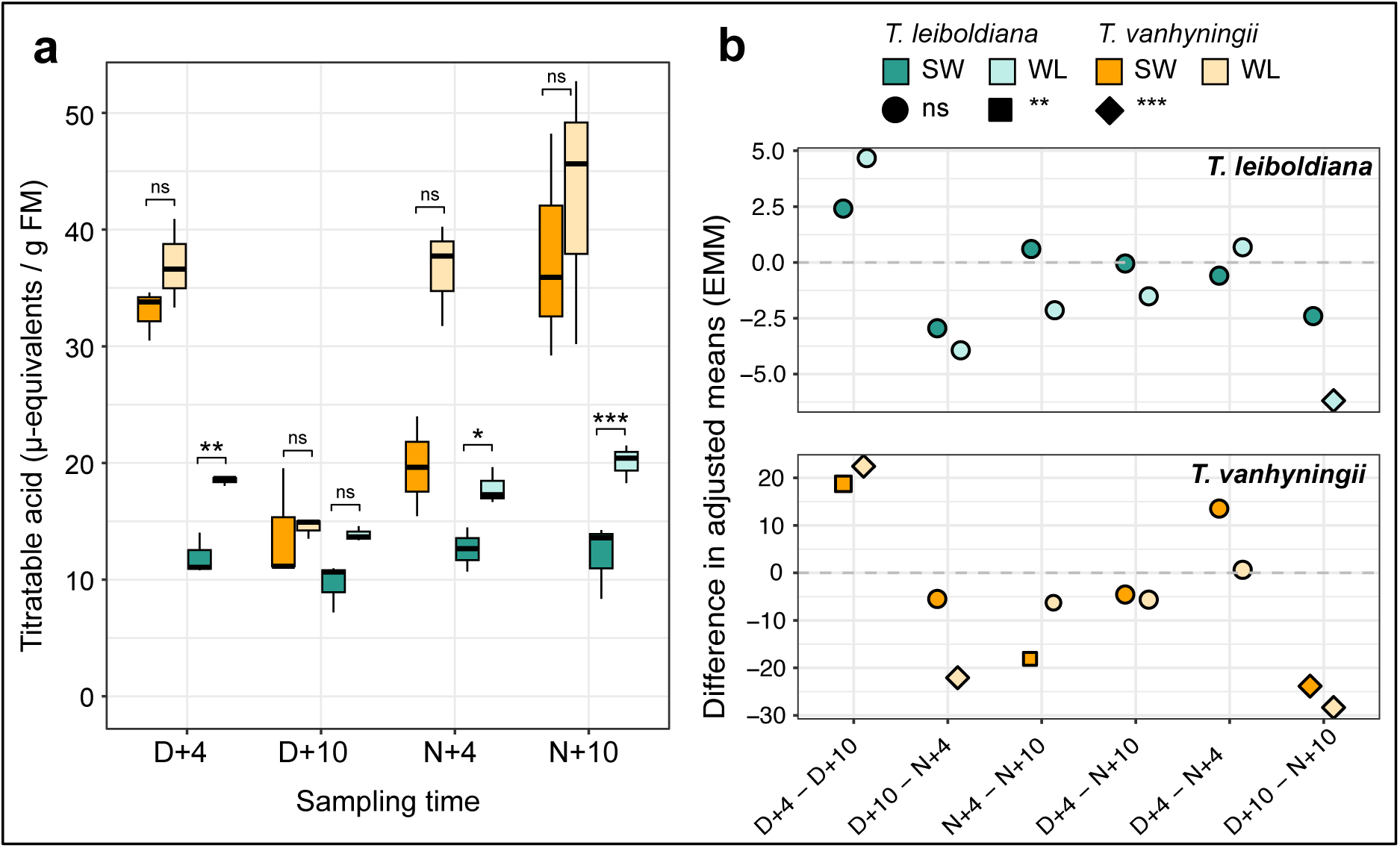
Titratable Acidity (TA) measurements of T. vanhyningii and T. leiboldiana. **a)** TA measured at four time points across a diel cycle under two watering regimes. The significance level of within-species, between-watering regime comparisons at a given time point from an LMM model are shown with following thresholds: ns - p-value > 0.05; * - p-value < 0.05; ** - p-value < 0.01; *** - p-value < 1E-03. **b)** Difference in adjusted means of TA values between each pair of timepoints, separated for each species. Each dot represents the difference between adjusted means of TA values between two timepoints within a species and watering regime, as estimated in the post-hoc Tukey HSD test. The first four timepoint comparisons are between adjacent timepoints that are 6 hours apart in a 24-hour cycle, whereas the last two represent differences in mean TA between timepoints that were 12 hours apart. The shape of each dot indicates the significance level of the difference in mean TA and is shown with the same thresholds as in a). Abbreviations: SW – standard watering conditions; WL – water-limited conditions; FM - Fresh Mass; D – Dawn; N – Nightfall.

### 4.2. Gas exchange in *T. leiboldiana*

At the start of the water-withholding experiment, *T. leiboldiana* showed a C_3_-like pattern of CO_2_ assimilation occurring principally during the day (Fig. 3a, Table S13). However, throughout day 2 and 3, net daytime CO_2_ uptake decreased in a step-wise manner till null through most of day 3. In days 1 to 3, there was no night-time net CO_2_ uptake. On day 4, a metabolic shift occurred towards a weak CAM-like pattern of night-time CO_2_ assimilation. From this point onwards, night-time net CO_2_ uptake became visible and increased gradually as the experiment progressed, with the highest values on day 8. *T. leiboldiana* also exhibited net CO_2_ uptake during the day, but this was consistently higher at night. After a 14-day recovery, the plant showed day-time net CO_2_ uptake and none during the night, an indication that it had fully shifted back to a C_3_-like photosynthetic metabolism (Fig. 3b, Table S14).

**Figure 3:**
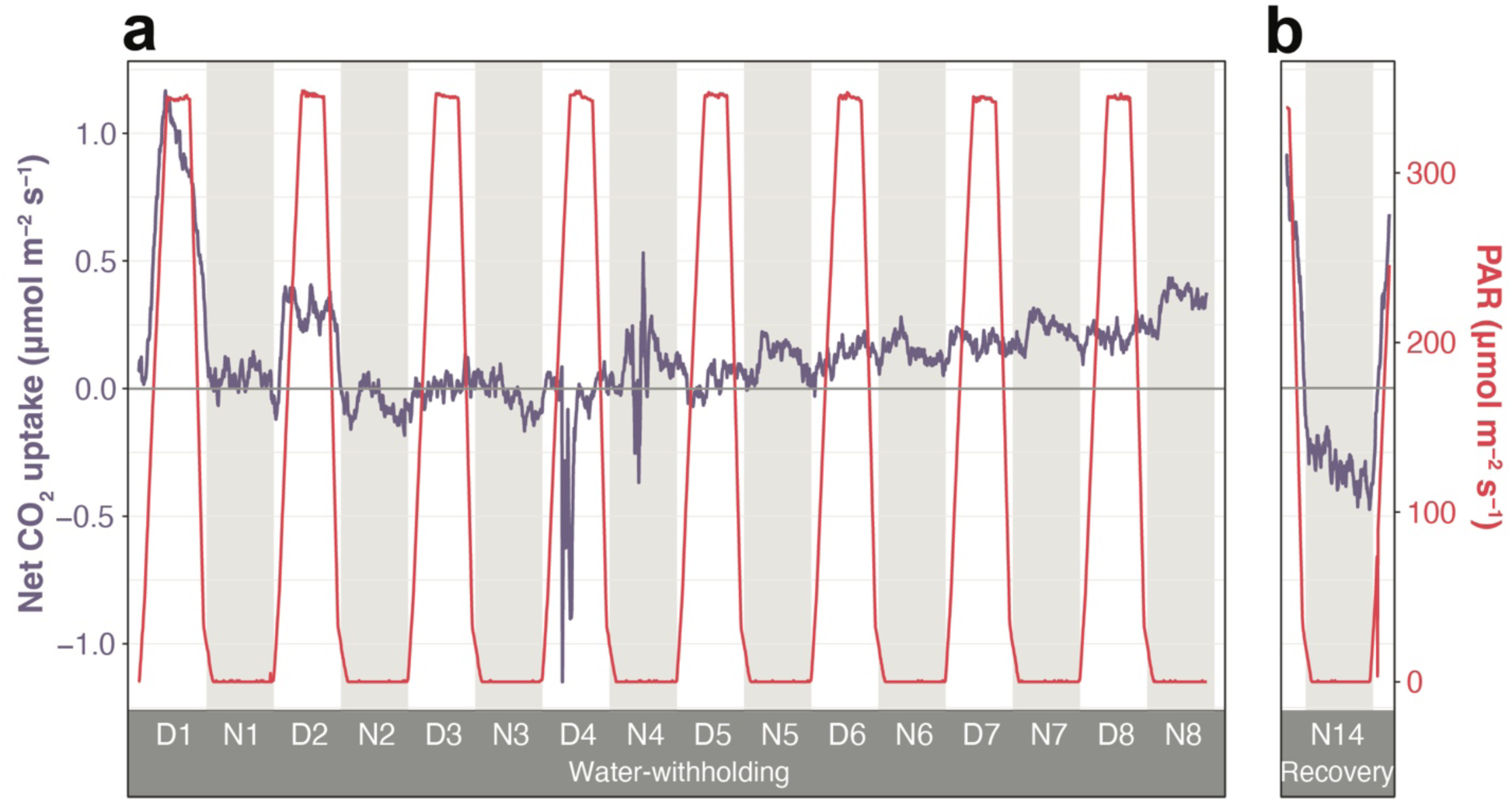
Gas exchange measurements of T. leiboldiana under a progression from well-watered to water-limited conditions, followed by recovery. Net CO_2_ uptake (blue line, left Y-axis) and Photosynthetically Active Radiation (PAR; red line, right Y-axis) in leaves of T. leiboldiana. For visualization purposes, exponential smoothing was applied to the net assimilation curves. When the value of net CO_2_ uptake is positive (above horizontal line), the plant is assimilating more CO_2_ than it is leaking CO_2_, the opposite is true for negative values. A 24-hour cycle is divided into a light period (white background) and a dark period (grey background). **a)** Measurements throughout an 8-day period without watering. **b)** Measurements across one 24-hour cycle after a 14-day recovery.

### 4.3. Differential gene expression

Unique mapping rates were high for both species, but *T. leiboldiana*’s reads mapped better to its own genome than *T. vanhyningii*’s reads to the *T. fasciculata* genome. (Fig. S2). On average, *T. leiboldiana* had 16.3% more uniquely assigned reads per sample than *T. vanhyningii* (Table S7). However, *T. vanhyningii* had 8.7% more genes with non-zero counts compared to *T. leiboldiana*. A comparison of downstream results when mapping *T. leiboldiana* to its own reference genome, versus to *T. fasciculata*, shows minimal effect (See supplementary note). *T. leiboldiana* is approximately twice as distantly related to *T. fasciculata* as *T. vanhyningii* is. Therefore, we expect a much smaller effect using *T. vanhyningii* alignments to *T. fasciculata* on the results presented in this study.

In *T. vanhyningii*, variance in read counts was largely explained by time point, while watering regime was partially resolved in the second component of the PCA (Fig. S3). In *T. leiboldiana*, the first PC separated samples by watering regime, and the second component showed a clear split by time point. Overall, *T. leiboldiana* showed clearer transcriptomic distinctions by time point and watering regime than *T. vanhyningii* (Fig. S3).

DGE analyses revealed widely different counts of DE genes along the time and watering regime axes. In both species, many genes were DE between day and night, regardless of watering regime (Fig. 4). Under SW, the number of up-regulated genes in *T. vanhyningii* was comparable between day and night, with only six more genes up-regulated at night than at day, but *T. leiboldiana* had 2.4x more genes up-regulated at day than at night. Under WL, more genes were up-regulated at night in both species compared to SW. While this pattern is noticeable in *T. vanhyningii*, with 57 % of DE genes up-regulated at night under WL compared to 50 % under SW, it is especially pronounced in *T. leiboldiana* (46 % of genes up-regulated at night under WL compared to 30 % under SW).

**Figure 4:**
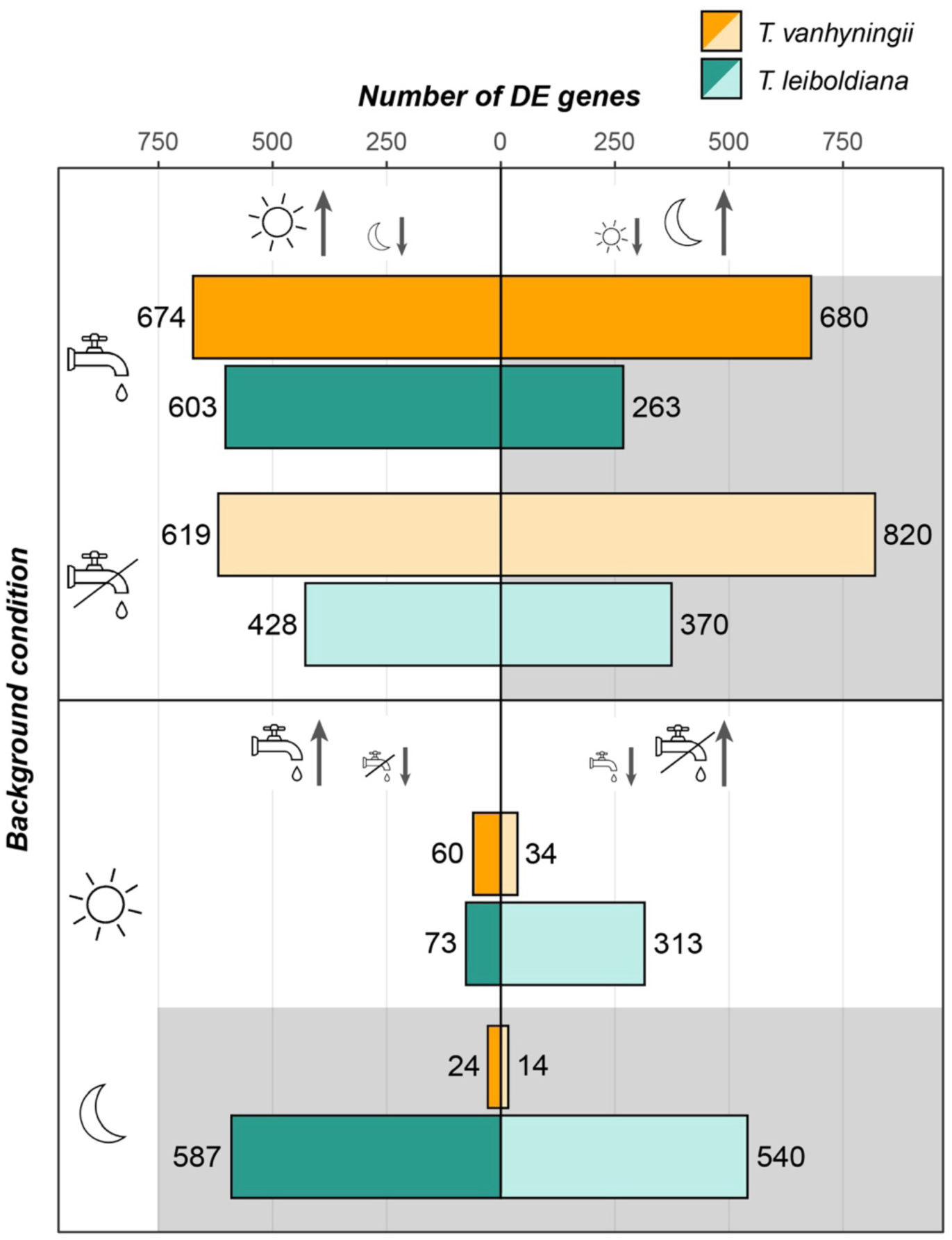
Counts of up- and down-regulated DE genes (FDR-adjusted p-value < 0.05, |Log(FC)| ≥ 1.5) in four comparisons across different background conditions. The first four bars show counts of time-dependent DE genes in two watering regimes, while the last four bars show counts of genes that are DE between watering regimes at two time points. Left-hand symbols indicate the background condition, e.g. the first two bars are counts of time-dependent DE genes under standard conditions. Left-hand bars show counts of genes that are up-regulated at day compared to night for time-dependent comparisons, and in water-limited compared to standard conditions for comparisons of watering regimes. Right-hand bars show counts of genes that are up-regulated at night compared to daytime for time-dependent comparisons, and in standard watering compared to water-limited conditions for comparisons of watering regimes. The DE analyses were conducted in each species separately. A dark-grey background indicates night-time, while lighter bar colours indicate water-limited conditions.

In comparison with time-dependent analyses, fewer genes were DE between watering regimes, except in *T. leiboldiana* at night. Contrary to *T. leiboldiana*, *T. vanhyningii* showed a very low number of DE genes between watering regimes, independent of time point, suggesting that most temporal expression patterns detected in the time-dependent DGE analysis were constitutive.

Compared to *T. vanhyningii*, *T. leiboldiana* manifested a more noticeable response to water withholding, especially at night. During the day, most genes appeared down-regulated under WL compared to SW (Fig. 4). The total number of DE genes between watering conditions at night was 2.9x larger than at day, which is a contrary trend to *T. vanhyningii*, where 2.5x more genes were DE between watering regimes at day than at night.

### 4.4. Overlap in transcriptomic response

The time-dependent transcriptomic response showed large heterogeneity across species and watering regimes, with 50 % of all orthogroups with one or more DE genes (hereafter DE orthogroups) being unique to one species and watering regime (Fig. 5a, sections a, b, c and d, Table S8). Only 5.9 % of all time-dependent DE orthogroups were shared between all groups (section abcd). These were enriched for photosynthesis and circadian clock-related functions (Table S9).

**Figure 5:**
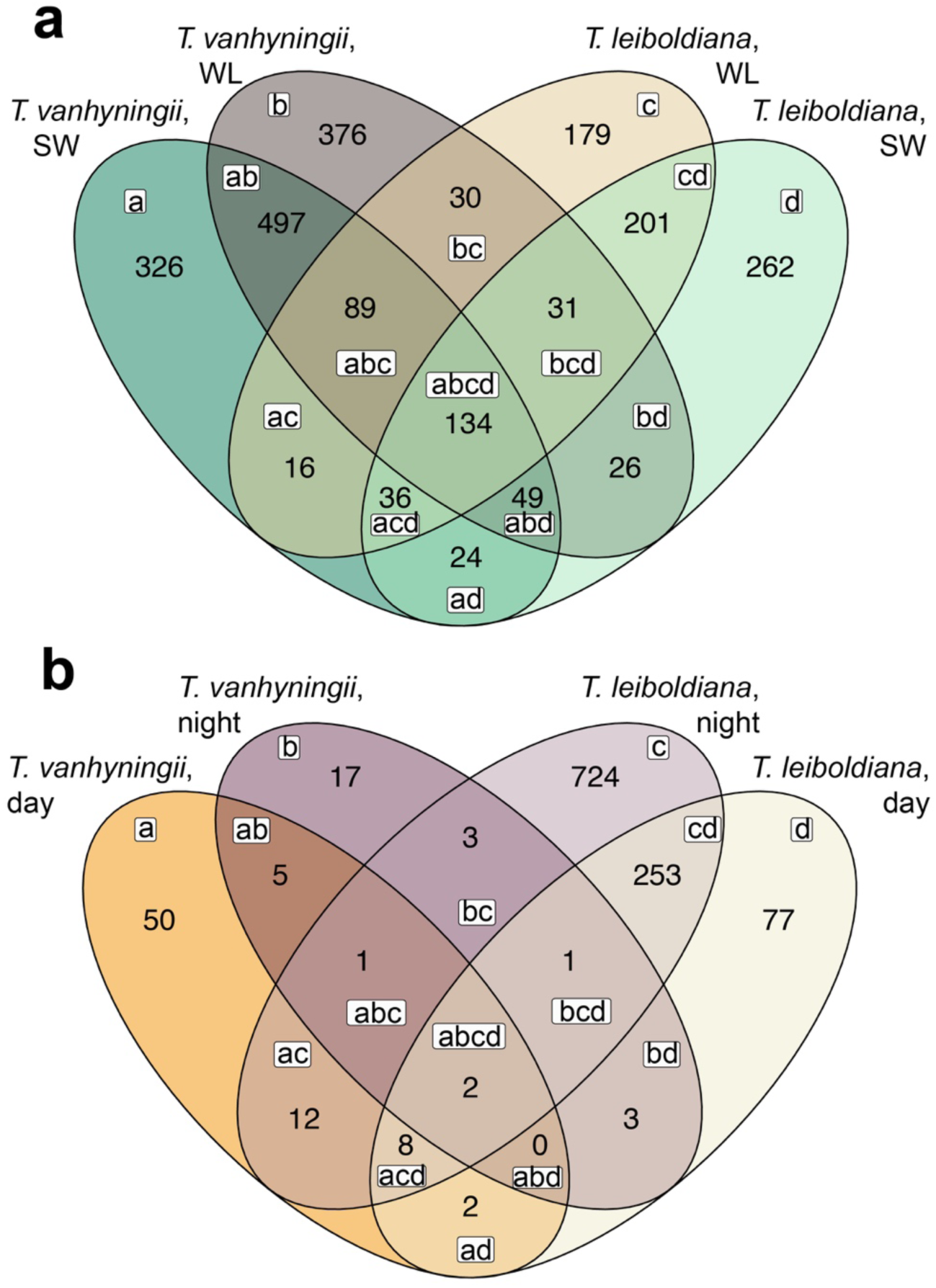
Overlap of DE orthogroups between species, time points and watering regimes. **a)** Venn diagram showing the number of shared time-dependent DE orthogroups (i.e., day versus night) per combined group of species and watering condition. Example: section a shows the number of orthogroups that contained genes that were exclusively DE between day and night under standard watering conditions in T. vanhyningii, while section ad shows the number of orthogroups with a gene that is DE both in T. leiboldiana and in T. vanhyningii under standard watering conditions. **b)** Venn diagram showing the number of shared DE orthogroups between watering regimes (i.e., water-limited versus standard conditions) per combined group of species and time point. Letters inside sections show the overlap between the four groups. Example: section a shows the number of orthogroups that contained genes that were exclusively DE between watering regimes at day-time in T. vanhyningii, while section ad shows the number of orthogroups with a gene that is DE both in T. leiboldiana and in T. vanhyningii at day-time.

The largest overlap was found within species, between conditions, as only 19 % of all time-dependent DE genes were shared between species (Table S8, intersections of a and b, and c and d). In *T. leiboldiana*, genes that were uniquely time-dependent DE under WL and not under SW (Fig. 5a, Table S9, sections d, bd, abd and ad) were enriched for responses to water deprivation, abiotic stress and abscisic acid (ABA), while genes that were only time-dependent DE under SW (Fig. 5a, Table S9, sections c, bc, abc and ac) lacked such enrichments, indicating that *T. leiboldiana* experienced drought-induced stress under WL (Table S9). Conversely, functions related to water deprivation, response to abiotic stress or ABA were enriched in time-dependent DE genes shared between watering regimes in *T. vanhyningii* (Fig. 5a, Table S9, sections ab, abc, abcd and abd).

Under WL, more time-dependent DE orthogroups were shared between species than under SW (17% vs. 14%, Table S8), with enriched functions related to stomatal movement, ABA, and starch metabolism (Fig. 5a, Table S9, sections ad, acd, and abd).

The response to watering regime was largely non-shared between groups, with 75 % of DE orthogroups being unique to one species and time point (Fig. 5b, sections a, b, c, and d) and only two DE orthogroups being shared among all groups (Fig. 5b, section abcd). *T. vanhyningii* showed little overlap between time points in the response to WL (8 %), while in *T. leiboldiana* 24 % of genes are DE between watering regimes at both time points (Table S8). Due to the small number of DE genes in *T. vanhyningii* between watering regimes, the overlap between species was very small (2.8 %).

In summary, both the overall transcriptomic response to day-night changes and to watering regimes showed little overlap between species, yet diel expression between species seemed to overlap slightly more under water-limited than under standard conditions, indicating an increase in the resemblance of the two species’ metabolism under water-limited conditions.

### 4.5. CAM-related transcriptomic overlap and differences between species, time points and conditions

#### 4.5.1. Overlap between species

In *T. leiboldiana*, CAM-related gene expression is likely to be associated both with differential expression between time points and between watering regimes, as it activated a CAM cycle under water-limited conditions. When considering all DE orthogroups summed up across species, time points and watering regimes, 516 contained genes that were both DE between time points and watering regimes across species.

Of these 516 orthogroups, the largest group was exclusively DE in *T. leiboldiana*, both between time points and watering regimes (Fig. S4), and included orthogroups encoding for aquaporin NIP5-1, an aluminium-activated malate transporter (ALMT) 12 and a sugar transporter 5. The second largest group consisted of 132 orthogroups (26 %) which displayed time-dependent expression in *T. vanhyningii* while being DE between watering regimes in *T. leiboldiana*, and included twelve previously described CAM-related orthogroups. Most notable were orthogroups encoding for the sugar and malate transporters SWEET14 and ALMT 9. *SWEET14* showed elevated expression under WL in both species, yet it seemed up-regulated at night in *T. leiboldiana,* while in *T. vanhyningii* its highest expression was during the day (Fig. 6). *ALMT 9* was more highly expressed in *T. leiboldiana* under SW, and showed lower expression under WL. *T. vanhyningii*’s expression of *ALMT 9*’s ortholog was comparatively lower to *T. leiboldiana*‘s across all time points and treatments (Fig. 6).

**Figure 6:**
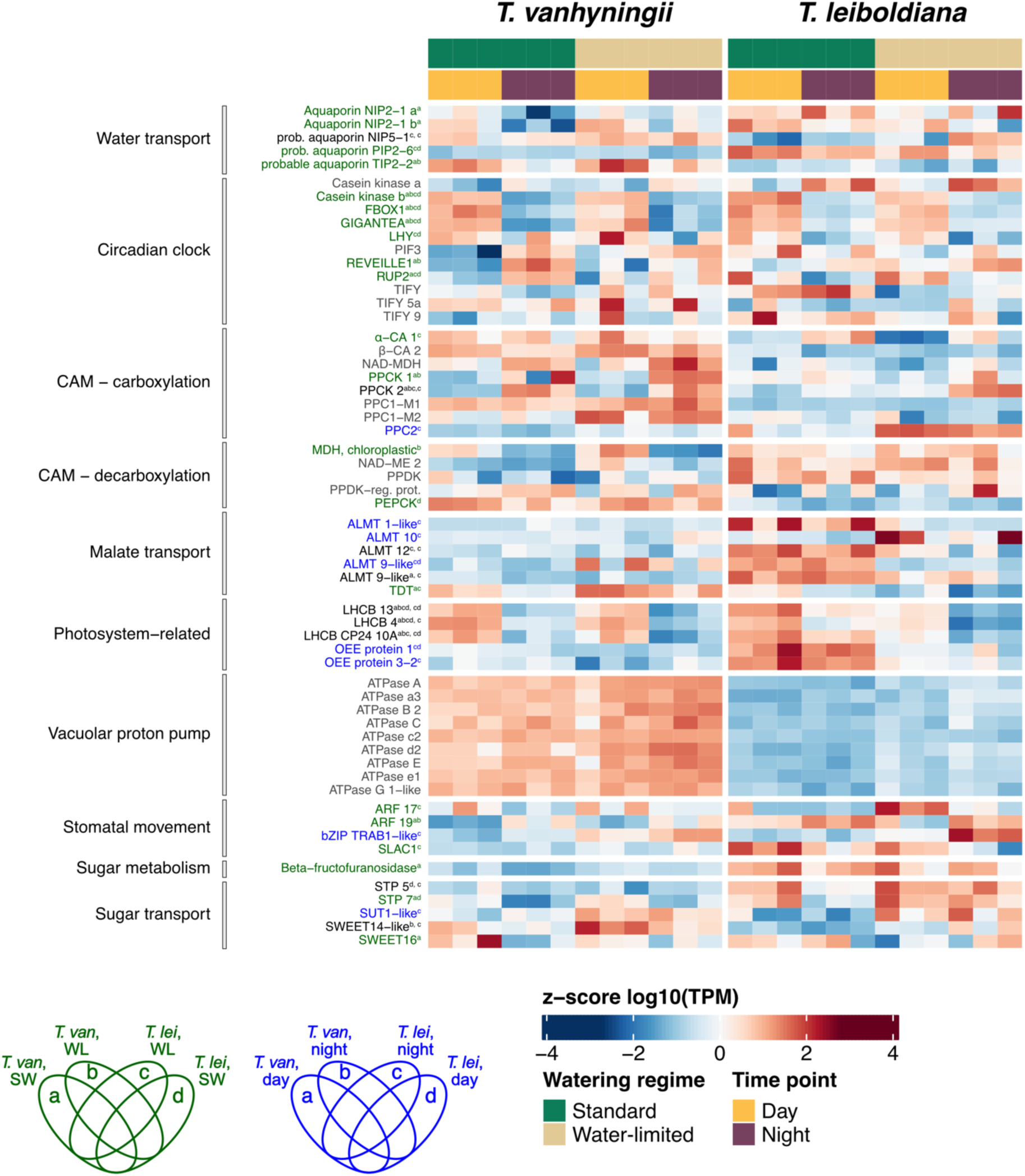
Transcript abundance in “Transcripts per Million” (TPM) for a subset of single-copy CAM-related orthogroups (listed in table S3), in T. vanhyningii (left panel, using the T. fasciculata ortholog) and T. leiboldiana (right panel, using the T. leiboldiana ortholog) at two time points and watering regimes. TPM values are transformed to a logarithmic scale and centred as a z-score. Displayed genes are organised in functional categories and gene name colour reflects their differential expression status: blue = DE between watering regimes, green = DE between time points, black is DE in both comparisons and grey = not DE. The letters in superscript indicate the intersection of the Venn diagram that each orthogroup belongs to. Schematics of Venn diagrams in the bottom left indicate what section each letter refers to. When an orthogroup is DE in both comparisons, the first superscripted letter indicates the intersection of the time-dependent Venn diagram (Fig. 5a), and the second superscripted letter, separated by a comma, indicates the intersection of the Venn diagram for watering regimes (Fig. 5b). Abbreviations: NIP - Nodulin 26-like Intrinsic Protein; PIP - Plasma Membrane Intrinsic Protein; TIP - Tonoplast Intrinsic Proteins; LHY - Late Elongated Hypocotyl; PIF - Phytochrome Interacting Factor; RUP - Repressor of UV-B Photomorphogenesis; α-CA - alpha Carbonic Anhydrase; β-CA - beta Carbonic Anhydrase; PPCK – Phosphoenolpyruvate carboxylase kinase; PPC – phosphoenolpyruvate carboxylase; NAD-MDH – Nad-dependent Malate Dehydrogenase; NAD-ME - NAD-dependent Malic Enzyme; PPDK - Pyruvate-Phosphate Dikinase; PPDK-reg. prot – PPDK-regulatory protein; PEPCK – Phosphoenolpyruvate Carboxykinase; LEA - Late Embryogenesis Abundant; ALMT – Aluminium-dependent Malate Transporter; TDT - Tonoplast Dicarboxylate Transporter; LHCB - Chlorophyll a-b binding protein; OEE - Oxygen Evolving Enhancer; ARF – Auxin Response Factor; bZIP - basic region leucine zipper; SLAC1 - slow-type anion channel 1; STP – Sugar Transporter Protein; SUT1 – Sucrose Transporter 1.

127 orthogroups (25 %) showed a time-dependent expression in both species but were only DE between watering conditions in *T. leiboldiana* (Fig. S4). Among these were 18 CAM-related orthogroups, yet only seven contained genes that were up-regulated under water-limited conditions in *T. leiboldiana*. This included an orthogroup encoding for PEPC kinase (*PPCK2*), which is crucial for CAM photosynthesis, as it activates the main carboxylating enzyme phosphoenolpyruvate carboxylase (PEPC, see section 4.4.4 for detailed discussion). Additionally, an orthogroup containing the drought resistance gene *YLS9* (Müller *et al*., 2017), and three orthogroups encoding for enzymes associated with glycolysis and gluconeogenesis were part of this group. *YLS9* showed time-dependent expression in *T. vanhyningii* in both watering regimes, while in *T. leiboldiana,* it acquired weakly time-dependent expression under WL (Fig. 6). Together with *PPCK2, YLS9* was one of the few CAM-related genes that exhibited an expression profile in *T. leiboldiana* that more closely resembled that of *T. vanhyningii* under WL.

We also searched for CAM-related orthogroups among the larger set of DE orthogroups called across time and watering regime separately. In *T. vanhyningii* and *T. leiboldiana*, 97 and 75 time-dependent DE CAM-related orthogroups were identified, respectively, revealing functions like circadian clock, glycolysis, stomatal movement, drought resistance, and water/sugar transport (Tables S5, S8). Overlap between species and watering regimes of these orthogroups followed a similar trend as the full set (51 % of genes unique to a species and watering regime). Twelve CAM-related DE orthogroups shared across all groups were linked to the circadian clock, photosynthesis, and photosystem activity (Table S10, section abcd). *T. vanhyningii* shared 46% of CAM DE genes between watering conditions, while *T. leiboldiana* shared 36%, indicating a stronger metabolic shift under WL. The proportion of time-dependent CAM DE orthogroups shared between species was slightly higher under WL (25 %) than under SW (22 %), yet remains a minority of the total set of CAM-related DE genes.

For DE orthogroups between watering regimes, six and 75 CAM-related orthogroups were present in *T. vanhyningii* and *T. leiboldiana* respectively (Table S8, Table S11). Only one orthogroup overlapped between species, whose orthologs encode for a probable linoleate 9S-lipoxygenase enzyme found differentially expressed between C_3_ and CAM *Tillandsia* species in a previous study (De La Harpe *et al*., 2020). Lipoxygenases are involved in the biosynthesis of jasmonates, which play an important role in response to various biotic and abiotic stressors (Bachmann *et al*., 2002).

#### 4.5.2. CAM-related expression in *Tillandsia vanhyningii*

In SW, *T. vanhyningii* showed time-dependent differential expression of core CAM gene *PPCK* and of genes underlying stomatal movement, vacuolar proton pumps, malate transport, sugar transport and glycolysis (Table S10). Several genes encoding for Late Embryogenesis Abundant (LEA) proteins linked to drought tolerance (Arumingtyas et al., 2013; Magwanga et al., 2018) also showed diel expression in SW.

46 CAM-related time-dependent DE genes overlapped between SW and WL (Table S8), including genes encoding for PPCK, circadian clock regulator REVEILLE1, a beta-amylase (starch breakdown), two auxin response factors (ARF, linked to stomatal function (Bouzroud *et al*., 2020)), two aquaporins and multiple LEA proteins (Table S10), indicating that genes from a wide set of CAM-related functions were expressed similarly between conditions. *AVP1*, a secondary proton pump that aids in the vacuolar transport of malate, was only DE in SW. A gene encoding for a chloroplastic malate dehydrogenase (MDH) showed diel expression in both conditions, though it was only significantly time-dependent under WL (Fig. 6). It was up-regulated during the day, suggesting a role in the day-time decarboxylation module of CAM. Its corresponding ortholog in *T. leiboldiana* showed no time-dependent expression in any condition. We also observed a shift in time-dependent expression between conditions among genes encoding for sugar transporters. While sugar transporter *SWEET16* showed significant time-dependent expression exclusively under SW, *SWEET14* was only DE under WL (Fig. 6). Both genes were up-regulated during the day when displaying time-dependent expression.

Very few CAM-related genes were DE in *T. vanhyningii* between watering regimes. They included several genes encoding enzymes associated with glycolysis and gluconeogenesis and genes previously described as DE between C_3_ and CAM *Tillandsia* (De La Harpe et al. 202). None of these genes were part of the core CAM pathway, and while genes encoding for many core CAM enzymes did show a slight increase in expression under water-limited conditions (β-carbonic anhydrase, NAD-dependent MDH, NAD-ME, Fig. 6), their expression was overall similar between watering regimes.

#### 4.5.3. CAM-related expression in *Tillandsia leiboldiana*

Under SW, time-dependent CAM-related DE genes were associated with functions that are typically expected to show temporal oscillation in plants that do not perform CAM (glycolysis and gluconeogenesis, sugar transport, circadian clock regulation). Under WL, 29 DE genes overlapped with standard conditions (Table S10), yet many additional CAM-related genes became time-dependent DE, involved in malate transport, stomatal movement, sugar metabolism, water transport and starch metabolism. Genes encoding Alpha-carbonic anhydrase 1 (α-CA) and phosphoenolpyruvate carboxylase kinase (PPCK) were up-regulated at night. *SLAC1* and two auxin response factors related to stomatal function were also time-dependent DE under WL (Table S10, Fig. 6). These findings further emphasise that a CAM cycle was activated in *T. leiboldiana* under WL.

Furthermore, most CAM-related genes that were DE between watering regimes could be found in *T. leiboldiana* at night (venn diagram section c) and included genes encoding the core CAM enzymes PEPC and PPCK (Table S11). Several genes encoding for malate transporters were also in this group, though they were generally down-regulated under WL, except for *ALMT10* (Fig. 6). Sugar transport genes were up-regulated under WL, such as genes encoding for sugar transporter proteins (STPs) and Sucrose Transporter 1 (*SUT1*), whose ortholog also showed elevated expression under WL in *T. vanhyningii*. Genes related to stomatal function were generally more highly expressed under WL but displayed varying behaviour in terms of the time of expression (Fig. 6). While *ARF17* and *SLAC1* were up-regulated during the day, *ARF19* and *TRAB1* were up-regulated during the night.

#### 4.5.4. Diel expression of PPC and PPCK homologs, and other CAM-related gene families, between species and watering regimes

The transcript abundances of PEPC and PPCK homologs revealed a variety of expression patterns. Gene *Tfasc_v1.16311-RA*, which is the ortholog of *PPC1-M1* (Fig. S5a) and was previously identified as the CAM-related fluctuating copy of PEPC in *T. fasciculata* (Groot Crego *et al*., 2024) did not show clear diel expression changes in *T. vanhyningii* (Fig. 7). In *T. leiboldiana* a slight increase was noticeable in its ortholog at night but was not significantly DE in either condition. However, *PEPC* was noticeably more highly expressed in *T. vanhyningii* compared to *T. leiboldiana*. The remaining known *PEPC* genes were generally less expressed than *PPC1-M1* in both species but appeared to become activated under WL, namely in *T. vanhyningii* for *PPC1-M2* (not significant), and in *T. leiboldiana* for *PPC2*.

**Figure 7:**
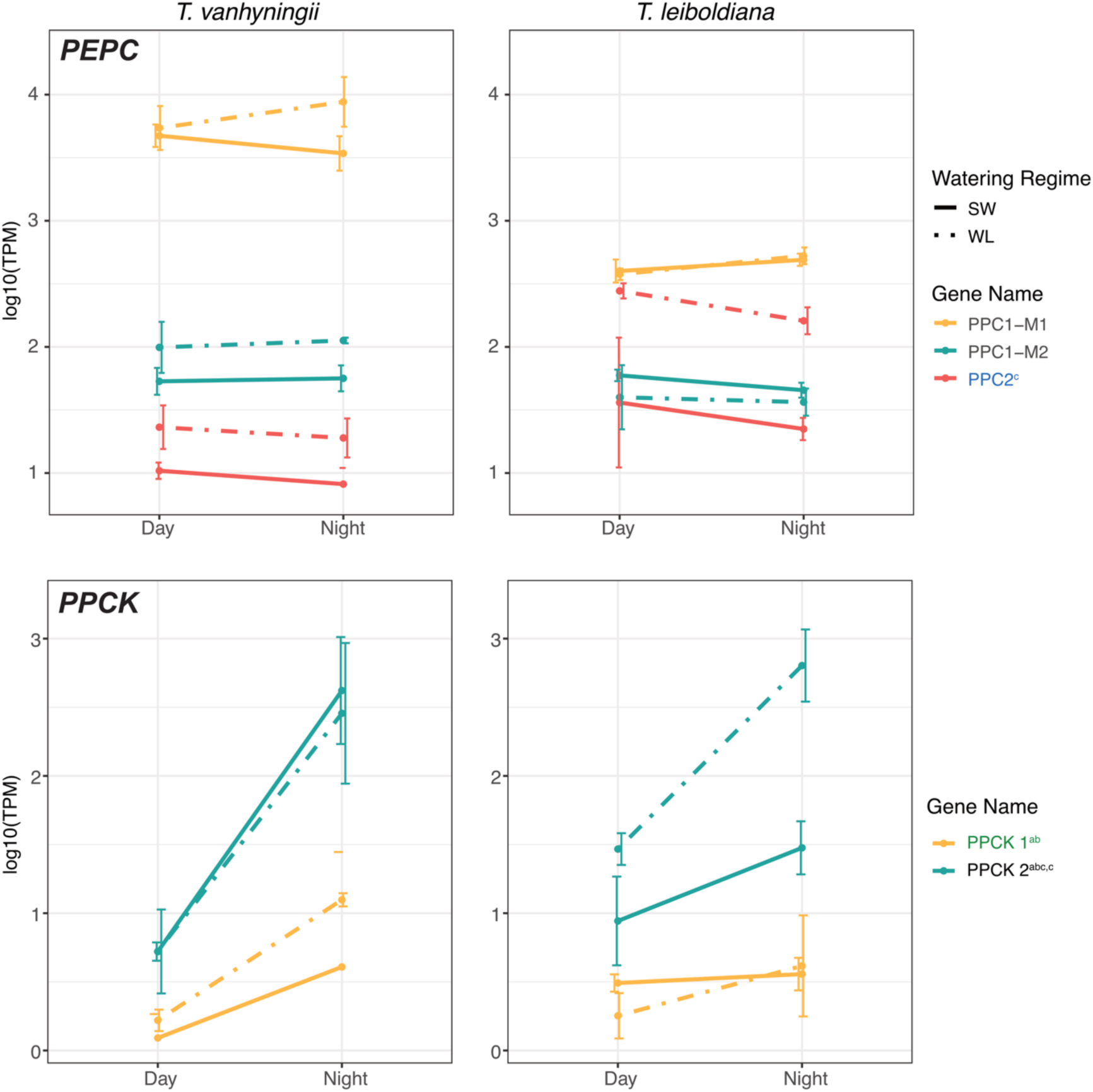
Transcript abundance in “Transcripts per Million” (TPM) for all homologous genes encoding for PEPC and PEPC kinase (PPCK) in T. vanhyningii (left panel) and T. leiboldiana (right panel) at two time points and watering regimes. TPM values are transformed to a logarithmic scale. The colour of gene names reflects their differential expression status: blue = DE between watering regimes, green = DE between time points, black is DE in both comparisons and grey = not DE. The superscripted letters after a gene name indicate the intersection of the Venn diagram this gene belonged to.

For PEPC kinase (PPCK), diel expression stood out in one gene (*PPCK2,* Fig. S6) in *T. vanhyningii* under both watering regimes. While its ortholog was not strongly expressed at night under SW in *T. leiboldiana*, it became highly expressed under WL (Fig. 7). *PPCK1* showed mild diel fluctuation in *T. vanhyningii* but its ortholog was overall lowly expressed in *T. leiboldiana*.

The NAD-dependent malate dehydrogenase (NAD-MDH, OG0001548), previously identified as CAM-related (Groot Crego et al., 2024), is crucial for night-time carboxylation in CAM. This gene has two copies in *A. comosus*, *T. fasciculata*, and *T. leiboldiana* (Fig. S7a). Neither copy showed significant differential expression in this study. One copy had increased night expression, while the other was higher during the day (Fig. S7b), with both showing slightly elevated expression under WL in both species. Despite similar time-dependent expression profiles, the constitutive CAM plant exhibited substantially higher expression of this and other genes encoding for core CAM enzymes, including V-ATPase, PEPCK, NAD-ME, PPDK, and PPDK-regulatory protein (Fig. 6, Fig. S8-S10).

### 4.6. Description of the general response to water withholding

GO enrichment of watering condition-dependent DE genes in *T. leiboldiana* revealed a pronounced drought response, with the top up-regulated genes linked to water deprivation and ABA signalling. Processes that were down-regulated under WL pointed at a general shutdown of metabolism, such as photosynthesis, transport, and biosynthetic processes (Fig. 8a, Table S12). In contrast, *T. vanhyningii* showed minimal drought-related responses, with no significant changes in regulation of photosynthesis or water deprivation genes. Yet, a few enriched GO terms suggested a limited drought response, including inositol, jasmonate, and molybdate metabolism (Fig. 8b, Table S12).

**Figure 8:**
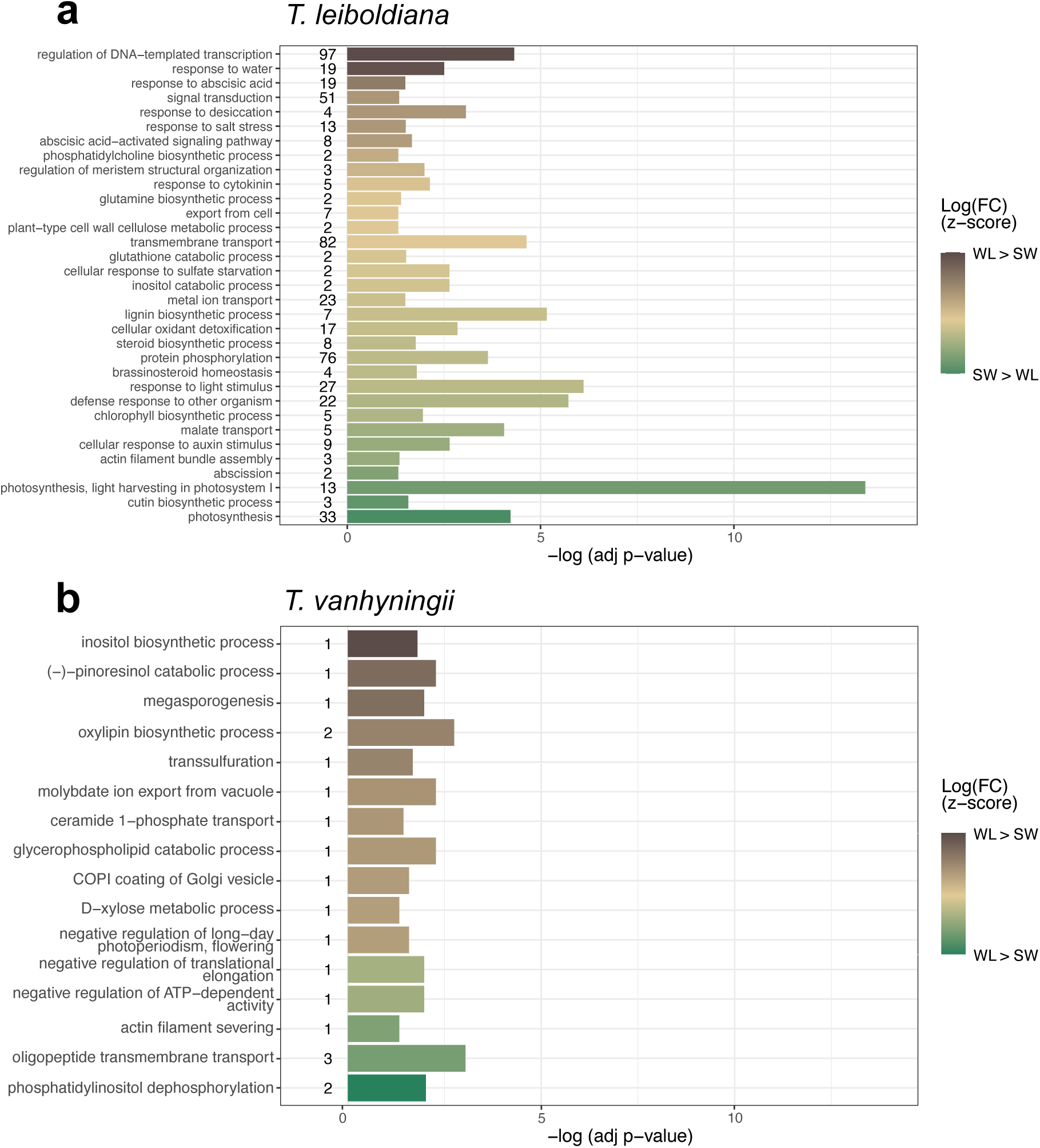
Summary of significantly enriched GO terms representing a Biological Process (BP) in DE genes between watering regimes in T. leiboldiana (**a**) and T. vanhyningii (**b**). GO term enrichment at day and night were combined for each bar plot. The bars show the significance of each enriched GO term as the p-value adjusted for multiple-testing, on a logarithmic scale. The colour gradient indicates up- and down-regulation of the genes underlying each GO term, as a z-score of the fold-change on a logarithmic scale. The GO terms are shown in descending order from most up-regulated in water-limited conditions to most up-regulated in standard conditions.

We detected 32 orthogroups related to drought response and water deprivation that were differentially expressed across species, watering regimes and time points. Eight of these were up-regulated under WL in *T. leiboldiana* compared to SW, and four were down-regulated (Fig. S12). The up-regulated orthogroups encoded for three LEA proteins, a phosphatase 2C, which is involved in ABA signalling (Jung *et al*., 2020), a galactinol-sucrose galactosyltransferase, which is associated with ROS scavenging (Nishizawa *et al*., 2008), an annexin which also confers oxidative protection under stressful conditions (Konopka-Postupolska *et al*., 2009), and the protein YLS9 (see section 4.4.1).

In *T. vanhyningii*, only five orthogroups related to drought-induced stress were differentially expressed between watering regimes, of which three were up-regulated under WL: an inositol-3-phosphate synthase, a 12-oxophytodienoate reductase involved in jasmonic acid biosynthesis and a molybdate ion transporter. In contrast to the low number of drought-related genes that were DE between watering regimes in *T. vanhyningii*, eighteen were DE between time points. However, out of these, sixteen already exhibited time-dependent expression under SW. Several orthologs also showed overall higher expression in *T. vanhyningii* than in *T. leiboldiana*. Genes encoding for LEA proteins 5 and 8, for example, were more highly expressed in *T. vanhyningii* and showed diel oscillation in both watering regimes, with their expression increasing at night (Fig. S12). Orthologous genes related to ABA signalling exhibited time-dependent expression in *T. vanhyningii* under both watering regimes (except for phosphatase 2C), and in some cases also in *T. leiboldiana* (Fig. S12). A 9-cis-epoxycarotenoid dioxygenase (NCED), involved in ABA biosynthesis (Wan *et al*., 2011), also showed diel expression changes in *T. vanhyningii* under SW. These findings suggest that high and/or diel expression of drought-related genes may be an inherent feature of constitutive CAM plants.

## 5. DISCUSSION

CAM is associated with diversification in many plant families and is considered a key innovation in Bromeliaceae (Benzing, 2000; Ogburn & Edwards, 2009; Quezada & Gianoli, 2011; Silvera et al., 2009). However, its evolutionary trajectory and the degree of genetic overlap between distinct CAM forms are not well understood (Edwards, 2023). The subgenus *Tillandsia* displays a wide spectrum of CAM phenotypes (Crayn et al., 2015; De La Harpe et al., 2020), but this diversity has so far been characterised only partially, mostly through carbon isotope ratios and transcriptomics under standard conditions. By comparing physiological and temporal transcriptomic responses to water limitation in two *Tillandsia* species, we aimed to explore the phenotypic range of CAM in this group and assess to what extent facultative and constitutive CAM rely on shared transcriptomic mechanisms.

### *Tillandsia leiboldiana* is a facultative CAM plant

Our TA, gas exchange and DGE analyses point at the presence of a latent CAM cycle in *T. leiboldiana* that is activated under drought-induced stress. First, water-withholding induced weak, yet significant night-time TA accumulation, absent in well-watered plants. Nocturnal acidification is a diagnostic feature of CAM (Winter & Smith, 2022) and has previously been used to identify facultative CAM in *Clusia* (Borland et al., 1992), *Mesembryanthemum* (Winter & Ziegler, 1992) and *Agave* (Heyduk et al., 2016).

Second, we observed a switch to net nocturnal CO_2_ uptake after four days of withholding water, followed by full recovery of day-time uptake after re-watering. Nocturnal net CO_2_ uptake is another hallmark of CAM, indicating open stomata and storage of CO_2_ in the vacuole in the form of malate (Lüttge, 1987), which aligns with our TA measurements. While the amplitude of nocturnal CO_2_ uptake was low, this pattern is comparable to other facultative CAM plants under drought. The CO_2_ uptake profiles in drought-stressed *Clusia pratensis*, *Calandrinia polyandra*, and *Talinum triangulare* all show modest night-time rates, and some daytime net assimilation remains (Winter & Holtum, 2014).

Transcriptomic signatures further support altered stomata behaviour for *T. leiboldiana*. Differential expression of stomatal function genes between watering regimes suggests altered stomatal behaviour under WL, though this may reflect a general drought-stress response rather than CAM-specific activity. For instance, TRAB1, involved in ABA signalling during drought stress (Hobo et al., 1999; Zhu, 2002), is up-regulated in *T. leiboldiana* under WL at night (Fig. 6). However, the guard cell anion channel SLAC1 controlling stomatal closure (Vahisalu *et al*., 2008) is down-regulated at night under WL (Fig. 6), potentially indicating that stomata open at night.

Third, night-time up-regulation of *PPCK2* in *T. leiboldiana* under WL indicates induction of the phosphorylation-based regulatory machinery necessary for nocturnal PEPC activity - a pattern that is entirely absent in SW.

Carbon isotope ratios reported for *T. leiboldiana* (δ^13^C = -28.0 to -31.3; Crayn et al., 2015; De La Harpe et al., 2020) would typically lead to its classification as a C_3_ plant. However, isotope values can mask facultative CAM activity, as seen in other bromeliads such as *Guzmania* and *Ronnbergia* (Pierce *et al*., 2002). Our integrative study therefore reveals that *T. leiboldiana* is not strictly C_3_, but a weak, facultative CAM species that performs C_3_ photosynthesis under benign conditions while acquiring weak CAM function under drought. Minimal expression changes in core CAM pathway genes between watering regimes further suggest that high transcript abundance of these genes may pre-condition facultative CAM expression in *Tillandsia*.

This raises an evolutionary question: what is the point of origin of CAM in the subgenus *Tillandsia*? Ancestral state reconstructions using carbon-isotope data support a C_3_ ancestor for the subgenus (De La Harpe *et al*., 2020), and multiple independent origins of CAM within the clade. However, the presence of inducible CAM in *T. leiboldiana* opens an alternative possibility - that at least parts of the CAM cycle may have been present prior to the divergence of the lineages leading to *T. leiboldiana* and *T. vanhyningii*. Although our pairwise design cannot rule out independent origins of facultative and constitutive CAM, it highlights the need for broader comparative datasets.

### *T. vanhyningii* shows minimal physiological and transcriptomic response to water withholding

Our physiological and transcriptomic analyses indicate that the quantitative response to water-withholding was far greater in *T. leiboldiana* than in *T. vanhyningii* (Figs. 2–4), suggesting that *T. leiboldiana*’s metabolism was affected by water deprivation, while *T. vanhyningii*’s seemed largely the same between conditions and therefore not substantially stressed. TA accumulated in a similar manner between watering regimes in *T. vanhyningii*, though accumulation occurred faster overnight under water-limited conditions. This aligns with studies on *T. ionantha*, which showed minimal physiological changes after six months of water withholding, attributed to daytime stomatal closure, low stomatal density, and efficient atmospheric moisture absorption via its dense trichome layer (E. J. Nowak & Martin, 1997; Ohrui et al., 2007).

A very small number of genes were found to be DE between watering regimes in *T. vanhyningii*, while a prominent response was detected in *T. leiboldiana*. Up-regulated enriched GO terms under WL in *T. vanhyningii* included metabolism of inositol, jasmonate and molybdate (Fig. 8b, Table S12), all known to play a role in drought-stress response (Li et al., 2022; Loewus & Murthy, 2000; Wasternack, 2014). However, the expression of many genes related to drought and abiotic stress response was time-dependent or more highly expressed in *T. vanhyningii* compared to *T. leiboldiana* under SW, including ABA biosynthesis and signalling genes, or LEA protein genes. A link between constitutive CAM phenotypes and temporal expression of ABA-related genes was observed by De La Harpe *et al*. (2020). ABA signalling plays an important role in stomatal closure as a response to water deprivation (Assmann & Jegla, 2016; Christmann et al., 2006; Wilkinson & Davies, 2002), but has also been associated with regulation of CAM photosynthesis (Mioto & Mercier, 2013; Ting, 1981). Our findings corroborate that time-dependent expression of ABA-related genes is characteristic of constitutive CAM phenotypes and does not necessarily indicate that *T. vanhyningii* was drought-stressed in our experiment.

Regarding the CAM-related response to WL, our findings showed that expression of CAM-related genes in *T. vanhyningii* remained largely similar. In some cases, however, such as sugar transporters SWEET16 and SWEET14, different genes performing the same functions were recruited for CAM under varying watering regimes.

Atmospheric or “grey” *Tillandsia* such as *T. vanhyningii* represent the most extreme form of adaptation to an epiphytic lifestyle among Bromeliaceae, having largely lost the ability to absorb water with roots (Benzing, 2000; Crayn *et al*., 2004). They occur in arid habitats and are all constitutive CAM plants, suggesting that CAM is a key innovation in this group (Crayn *et al*., 2004). Our findings confirm that atmospheric Tillandsia are drought-tolerant, as previously shown physiologically (Castillo et al., 2016; Martin, 1994; M. A. Nowak et al., 1997; Stiles & Martin, 1996), but now supported at the transcriptomic level, as *T. vanhyningii* was only mildly affected by water-limited conditions. However, while this could be attributable to *T. vanhyningii*’s constitutive CAM phenotype, other typical characteristics of atmospheric *Tillandsia* such as a dense trichome layer, need to also be considered as important contributors to their drought-tolerant nature.

### Shared versus non-shared transcriptional architecture of photosynthetic metabolism

Despite the relatively recent divergence of *T. vanhyningii* and *T. leiboldiana* (Barfuss et al., 2016; Mendoza et al., 2017; Yardeni et al., 2025), the two species show clear physiological and transcriptomic differentiation consistent with distinct photosynthetic phenotypes. Only limited overlap in time-dependent and watering-dependent DE genes was found between species, mirroring observations in other CAM lineages such as Agavoideae (Heyduk *et al*., 2022).

Most CAM-related, time-dependent DE genes showed diel expression cycling in only one species (Fig. 9). Although *T. leiboldiana* seemingly activated CAM under WL, the underlying transcriptomic architecture differed substantially from the constitutive CAM pattern of *T. vanhyningii*. Notable exceptions were *PPCK2* (Fig. 7, Fig. 9) and the drought-associated protein YLS9, which showed a similar time-dependent expression both in *T. vanhyningii* and drought-stressed *T. leiboldiana*. This restricted overlap suggests that the facultative and constitutive CAM cycles of these species rely on largely distinct transcriptional solutions, rather than expression-level modulation of a common pathway.

**Figure 9:**
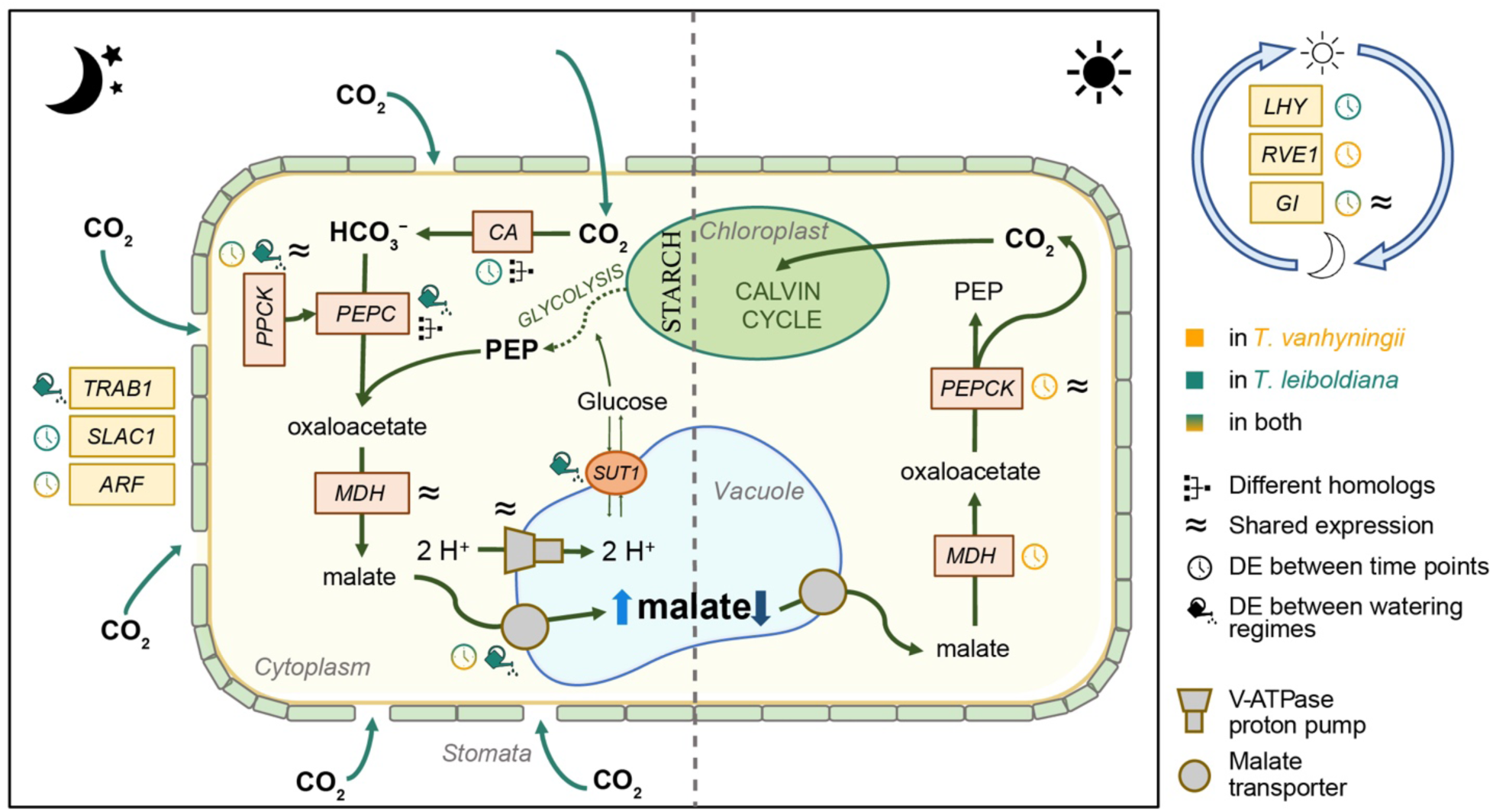
Schematic of the CAM pathway (adapted from Groot Crego et al. 2024), with an overview of CAM-related gene expression in standard watering and water-limiting conditions in T. vanhyningii and T. leiboldiana. Enzymes are shown in boxes, while pathway products are bold outside boxes. Enzymatic members of CAM metabolism pathways are shown in orange, while stomatal and circadian regulators are highlighted in yellow. Stomatal regulators are on the left outside the cell and circadian regulators on the right. Symbols indicate whether genes were DE in this study in timewise comparisons (clock symbol) or between watering regimes (watering can symbol). The color of the symbol indicates in which species the differential expression was detected, while a gradient of both colors indicates DE in both. “Different homologs” indicates that the species show CAM-like expression in distinct homologs of a gene family underlying the enzyme. “Shared expression” indicates high similarity in expression pattern between species across conditions. Abbreviations: CA – Carbonic Anhydrase, PEPC – Phosphoenolpyruvate Carboxylase, PPCK – Phosphoenolpyruvate Carboxylase Kinase, MDH – Malate Dehydrogenase, PEPCK - Phosphoenolpyruvate Carboxykinase, SLAC1 - slow-type anion channel 1, ARF - Auxin Response Factor, SUT1 – Sucrose Transporter 1, LHY - Late Elongated Hypocotyl, RVE1 – REVEILLE 1, GI – protein GIGANTEA.

This is supported by the distinct expression patterns of gene family members of PEPC, a core CAM gene. Under WL, the homologous copy of *PPC2* appeared up-regulated in *T. leiboldiana*, while being lowly expressed under all conditions in *T. vanhyningii*. Because none of the PEPC copies are time-dependently expressed in this experiment, likely due to the time of sampling not being synchronous with the peak expression times of these genes, it is difficult to confirm that *PPC2* is active in CAM in *T. leiboldiana* under drought-stress. Both in *T. fasciculata* and in *Agave*, PEPC gene expression peaks at dusk, while samples were taken 4 hours after dusk in this study (Abraham et al., 2016; Groot Crego et al., 2024). While in the majority of CAM plant lineages, including *T. fasciculata* (Groot Crego *et al*., 2024) and *A. comosus* (Ming *et al*., 2015), the PEPC gene copy recruited into the CAM pathway is *PPC1*, recruitment of *PPC2* has been suggested in *Yucca aloifolia* (Heyduk *et al*., 2022) and in *Isoetes taiwanensis* (Wickell *et al*., 2021). This supports the view that PEPC paralog recruitment can evolve flexibly within CAM lineages.

Another core CAM gene that showed increased time-dependent expression in *T. leiboldiana* under WL was α-carbonic anhydrase (α-CA, Fig. S11). Carbonic anhydrases convert CO_2_ to HCO_3_^-^, the first step of carbon assimilation in CAM photosynthesis. In pineapple, however, β-CA, rather than α-CA are modulated in CAM (Ming *et al*., 2015), and *T. vanhyningii* does not share the time-dependent expression of this gene. In several *Yucca* species, on the other hand, α-CA did show time-dependent expression under water-limited conditions (Heyduk *et al*., 2019). The CAM-like expression of *PPC2* and α-CA in *T. leiboldiana* suggests species-specific recruitment of different genes and homologs into the CAM pathway (Fig. 9).

Except for PPCK*, PPC2* and α-CA, the core (de)carboxylation module seems relatively conserved between species. However, other CAM-associated processes (Fig. 9), such as malate transport, sugar transport, stomatal regulation, water channels and starch metabolism, exhibited substantial transcriptomic divergence between species. Overall, the realized CAM phenotypes are underpinned by a complex mixture of conserved patterns at core genes and lineage-specific changes at peripheral genes.

## Conclusions

This study shows that one of the most C_3_-like members of subgenus *Tillandsia*, *T. leiboldiana*, is not fully C_3_, but a weak, facultative CAM plant. In contrast, *T. vanhyningii* exhibited strong physiological and transcriptomic drought tolerance consistent with its constitutive CAM phenotype and epiphytic lifestyle. Although these closely related species accumulated organic acids overnight under water limitation, they deployed largely distinct transcriptional architectures, particularly outside the core CAM (de)carboxylation module. This suggests that the evolutionary trajectory linking facultative and constitutive CAM involves broad transcriptional rewiring at peripherical parts of the networks, not simply the quantitative up-regulation of a conserved gene set. The remarkable phenotypic and transcriptomic diversity within the subgenus *Tillandsia* highlights its potential as an outstanding system for exploring the CAM continuum and the evolutionary flexibility underlying this key innovation.

## 8. ACKNOWLEDGEMENTS

We thank Viktor Vagovic at the Core Facility Botanical Garden of the University of Vienna for caretaking and maintenance of the plant accessions used in this study and assistance with the drought experiment. We also thank Aglaia Szukala, Marta Pelizzola, Nina Flowers and Simon Reynaert for advice and discussion on transcriptome analysis. We thank Antony Crisp for advice and discussions on titration assays.

## 10. DATA AVAILABILITY AND BENEFIT-SHARING

TA and gas exchange measurements, detailed information on statistical testing and accession information is provided in the supplementary tables of this manuscript. The raw RNA-seq data is available at the NCBI SRA database under the BioProject <u>PRJNA1121978</u>. Scripts, read counts and extended tables of GO enrichment analyses are available on the following repository: https://github.com/cgrootcrego/Tillandsia_CAM-Drought.

Benefits from this research accrue from the public dissemination of our data and findings through open access repositories, as detailed above.

## 11. AUTHOR CONTRIBUTIONS

This study was conceived by CL, JH, MDH, OP, and CGC. Accessions were sampled by WT. The greenhouse experiment and sampling were conducted by MDH and MHJB. Laboratory work including TA measurements and RNA extraction and library prep was conducted by MDH and MHJB under the guidance of OP. Analyses were performed by CGC, MDH and SS. Gas exchange measurements were done by GB with input from WW. The manuscript was written by CGC with initial contributions from SS, was amended by OP, and was finally read and approved by all coauthors.

## 12. SUPPLEMENTARY DATA - CAPTIONS

Figure S1: Diagnostic plots for Linear Mixed-Effects Models constructed in each species to assess the effect of time and watering regime on titratable acidity measurements.

Figure S2: Mapping rates across 24 samples of different species and watering regimes.

Figure S3: Principal component analysis of TMM-normalised read counts in log(CPM).

Figure S4: Overview of the overlap between DE analyses across time points and across watering regimes.

Figure S5: Gene trees of PPC1 (A) and PPC2 (B).

Figure S6: Gene trees of PPCK with the isozymes PPCK1 and PPCK2.

Figure S7: Expression of MDH orthologs.

Figure S8: Expression of NAD-ME.

Figure S9: Expression of PEPCK.

Figure S10: Expression of PPDK and PPDK regulatory protein.

Figure S11: Expression of carbonic anhydrases.

Figure S12: Transcript abundance in “Transcripts per Million” (TPM) for a subset of single-copy orthogroups related to drought response in *T. vanhyningii* and *T. leiboldiana*

Table S1: Sampling overview and summary of RNA extractions for 24 samples.

Table S2: Full list of CAM-related orthogroups obtained from Yardeni et al. 2021 based on A. comosus (pineapple) genes.

Table S3: List of 61 CAM-related orthogroups selected for expression visualization in Figure 4.

Table S4: List of 31 selected orthogroups related to water deprivation and drought response for expression visualization in Figure S7.

Table S5: Titratable acidity measurements for Tillandsia vanhyningii and Tillandsia leiboldiana at four time points spread over 6-hour intervals.

Table S6: Statistical models and results for titratable acidity measurements. Table S7: Summary of per-gene read counting with FeatureCounts.

Table S8: Overlap of DE orthogroups between species, watering regimes and time points.

Table S9: List of significantly enriched GO terms in each intersection of the Venn diagram showing overlap of DE genes between time points.

Table S10: Description of CAM genes in sections of the Venn diagram A (Day vs Night).

Table S11: Description of CAM genes in sections of the Venn diagram B (Water-limited vs standard watering conditions).

Table S12: List of significantly enriched GO terms in each intersection of the Venn diagram showing overlap of DE genes between watering regimes.

Table S13: Gas exchange measurements of *T. leiboldiana* under a progression from well-watered to water-limited conditions.

Table S14: Gas exchange measurements of *T. leiboldiana* after a 14 days recovery following water-withholding.

## REFERENCES

Abraham, P. E., Yin, H., Borland, A. M., Weighill, D., Lim, S. D., De Paoli, H. C., Engle, N., Jones, P. C., Agh, R., Weston, D. J., Wullschleger, S. D., Tschaplinski, T., Jacobson, D., Cushman, J. C., Hettich, R. L., Tuskan, G. A., & Yang, X. (2016). Transcript, protein and metabolite temporal dynamics in the CAM plant Agave. Nature Plants, 2(12), 16178.

Alexa, A., Rahnenführer, J., & Lengauer, T. (2006). Improved scoring of functional groups from gene expression data by decorrelating GO graph structure. Bioinformatics, 22(13), 1600–1607.

Arumingtyas, E. L., Savitri, E. S., & Purwoningrahayu, R. D. (2013). Protein profiles and dehydrin accumulation in some soybean varieties (*Glycine max* L. merr) in drought stress conditions. American Journal of Plant Sciences, 04(01), 134–141.

Assmann, S. M., & Jegla, T. (2016). Guard cell sensory systems: recent insights on stomatal responses to light, abscisic acid, and CO2. Current Opinion in Plant Biology, 33, 157–167.

Bachmann, A., Hause, B., Maucher, H., Garbe, E., Vörös, K., Weichert, H., Wasternack, C., & Feussner, I. (2002). Jasmonate-induced lipid peroxidation in barley leaves initiated by distinct 13-LOX forms of chloroplasts. Biological Chemistry, 383(10), 1645–1657.

Barfuss, M. H. J., Till, W., Leme, E. M. C., Pinzón, J. P., Manzanares, J. M., Halbritter, H., Samuel, R., & Brown, G. K. (2016). Taxonomic revision of bromeliaceae subfam. Tillandsioideae based on a multi-locus DNA sequence phylogeny and morphology. Phytotaxa, 279(1), 1–97.

Barghi, N., Hermisson, J., & Schlötterer, C. (2020). Polygenic adaptation: a unifying framework to understand positive selection. Nature Reviews. Genetics, 21(12), 769–781.

Barghi, N., Tobler, R., Nolte, V., Jakšić, A. M., Mallard, F., Otte, K. A., Dolezal, M., Taus, T., Kofler, R., & Schlötterer, C. (2019). Genetic redundancy fuels polygenic adaptation in Drosophila. PLoS Biology, 17(2), e3000128.

Bates, D., Mächler, M., Bolker, B., & Walker, S. (2015). Fitting Linear Mixed-Effects Models Using lme4. Journal of Statistical Software, 67, 1–48.

Benzing, D. H. (2000). Bromeliaceae: Profile of an Adaptive Radiation. Cambridge University Press.

Beutelspacher, C. R., & García-Martínez, R. (2021). Descripción de nuevos híbridos y una nueva combinación en Tillandsia (Bromeliaceae) para Chiapas, México. Lacandonia, 15(2), 19–28.

Borland, A. M., Griffiths, H., Maxwell, C., Broadmeadow, M. S. J., Griffiths, N. M., & Barnes, J. D. (1992). On the ecophysiology of the Clusiaceae in Trinidad: expression of CAM in Clusia minor L. during the transition from wet to dry season and characterization of three endemic species. The New Phytologist, 122(2), 349–357.

Borland, A. M., Guo, H.-B., Yang, X., & Cushman, J. C. (2016). Orchestration of carbohydrate processing for crassulacean acid metabolism. Current Opinion in Plant Biology, 31, 118–124.

Borland, A. M., Hartwell, J., Weston, D. J., Schlauch, K. A., Tschaplinski, T. J., Tuskan, G. A., Yang, X., & Cushman, J. C. (2014). Engineering crassulacean acid metabolism to improve water-use efficiency. Trends in Plant Science, 19(5), 327–338.

Bouzroud, S., Gasparini, K., Hu, G., Barbosa, M. A. M., Rosa, B. L., Fahr, M., Bendaou, N., Bouzayen, M., Zsögön, A., Smouni, A., & Zouine, M. (2020). Down regulation and loss of auxin response factor 4 function using CRISPR/Cas9 alters plant growth, stomatal function and improves tomato tolerance to salinity and osmotic stress. Genes, 11(3), 272.

Bräutigam, A., Schlüter, U., Eisenhut, M., & Gowik, U. (2017). On the Evolutionary Origin of CAM Photosynthesis. Plant Physiology, 174(2), 473–477.

Castillo, R. J., Cervera, J. C., & Navarro-Alberto, J. (2016). Drought and extreme temperature tolerance for Tillandsia dasyliriifolia, an epiphytic bromeliad from the northern coastal dune scrubland in Yucatan, Mexico. Botanical Sciences, 94(1), 121–126.

Chen, H., & Boutros, P. C. (2011). VennDiagram: a package for the generation of highly-customizable Venn and Euler diagrams in R. BMC Bioinformatics, 12, 35.

Christmann, A., Moes, D., Himmelbach, A., Yang, Y., Tang, Y., & Grill, E. (2006). Integration of abscisic acid signalling into plant responses. *Plant Biology (Stuttgart*, Germany), 8(3), 314–325.

Conway, J. R., Lex, A., & Gehlenborg, N. (2017). UpSetR: an R package for the visualization of intersecting sets and their properties. BioRxiv, 33, 2938–2940.

Crayn, D. M., Winter, K., Schulte, K., & Smith, J. A. C. (2015). Photosynthetic pathways in Bromeliaceae: Phylogenetic and ecological significance of CAM and C3 based on carbon isotope ratios for 1893 species. Botanical Journal of the Linnean Society. Linnean Society of London, 178(2), 169–221.

Crayn, D. M., Winter, K., & Smith, J. A. C. (2004). Multiple origins of crassulacean acid metabolism and the epiphytic habit in the Neotropical family Bromeliaceae. Proceedings of the National Academy of Sciences, 101(10), 3703–3708.

De La Harpe, M., Paris, M., Hess, J., Barfuss, M. H. J., Serrano-Serrano, M. L., Ghatak, A., Chaturvedi, P., Weckwerth, W., Till, W., Salamin, N., Wai, C. M., Ming, R., & Lexer, C. (2020). Genomic footprints of repeated evolution of CAM photosynthesis in a Neotropical species radiation. Plant, Cell & Environment, 43(12), 2987–3001.

Deng, H., Zhang, L.-S., Zhang, G.-Q., Zheng, B.-Q., Liu, Z.-J., & Wang, Y. (2016). Evolutionary history of PEPC genes in green plants: Implications for the evolution of CAM in orchids. Molecular Phylogenetics and Evolution, 94(Pt B), 559–564.

Dobin, A., Davis, C. A., Schlesinger, F., Drenkow, J., Zaleski, C., Jha, S., Batut, P., Chaisson, M., & Gingeras, T. R. (2013). STAR: Ultrafast universal RNA-seq aligner. Bioinformatics, 29(1), 15–21.

Edwards, E. J. (2023). Reconciling continuous and discrete models of C4 and CAM evolution. Annals of Botany. 10.1093/aob/mcad125

Ewels, P., Magnusson, M., Lundin, S., & Käller, M. (2016). MultiQC: summarize analysis results for multiple tools and samples in a single report. Bioinformatics, 32(19), 3047–3048.

Gibson, A. C. (1982). The anatomy of succulence. Crassulacean Acid Metabolism, 1–17.

Gilman, I. S., Smith, J. A. C., Holtum, J. A. M., Sage, R. F., Silvera, K., Winter, K., & Edwards, E. J. (2023). The CAM lineages of planet Earth. Annals of Botany, 132(4), 627–654.

Givnish, T. J., Barfuss, M. H. J., Van Ee, B., Riina, R., Schulte, K., Horres, R., Gonsiska, P. A., Jabaily, R. S., Crayn, D. M., Smith, J. A. C., Winter, K., Brown, G. K., Evans, T. M., Holst, B. K., Luther, H., Till, W., Zizka, G., Berry, P. E., & Sytsma, K. J. (2014). Adaptive radiation, correlated and contingent evolution, and net species diversification in Bromeliaceae. Molecular Phylogenetics and Evolution, 71(1), 55–78.

Goldstein, D. B., & Holsinger, K. E. (1992). Maintenance of polygenic variation in spatially structured populations: roles for local mating and genetic redundancy. Evolution; International Journal of Organic Evolution, 46(2), 412–429.

Griffiths, H., Robe, W. E., Girnus, J., & Maxwell, K. (2008). Leaf succulence determines the interplay between carboxylase systems and light use during Crassulacean acid metabolism in Kalanchoë species*. Journal of Experimental Botany, 59(7), 1851–1861.

Groot Crego, C., Hess, J., Yardeni, G., de La Harpe, M., Priemer, C., Beclin, F., Saadain, S., Cauz-Santos, L. A., Temsch, E. M., Weiss-Schneeweiss, H., Barfuss, M. H. J., Till, W., Weckwerth, W., Heyduk, K., Lexer, C., Paun, O., & Leroy, T. (2024). CAM evolution is associated with gene family expansion in an explosive bromeliad radiation. The Plant Cell. 10.1093/plcell/koae130

Gu, Z., Eils, R., & Schlesner, M. (2016). Complex heatmaps reveal patterns and correlations in multidimensional genomic data. Bioinformatics, 32(18), 2847–2849.

Hämälä, T., Guiltinan, M. J., Marden, J. H., Maximova, S. N., dePamphilis, C. W., & Tiffin, P. (2020). Gene Expression Modularity Reveals Footprints of Polygenic Adaptation in Theobroma cacao. Molecular Biology and Evolution, 37(1), 110–123.

Heyduk, K., Burrell, N., Lalani, F., & Leebens-Mack, J. (2016). Gas exchange and leaf anatomy of a C3-CAM hybrid, Yucca gloriosa (Asparagaceae). Journal of Experimental Botany, 67(5), 1369–1379.

Heyduk, K., McAssey, E. V., & Leebens-Mack, J. (2022). Differential timing of gene expression and recruitment in independent origins of CAM in the Agavoideae (Asparagaceae). The New Phytologist, 235(5), 2111–2126.

Heyduk, K., Ray, J. N., Ayyampalayam, S., & Leebens-Mack, J. (2018). Shifts in gene expression profiles are associated with weak and strong Crassulacean acid metabolism. American Journal of Botany, 105(3), 587–601.

Heyduk, K., Ray, J. N., Ayyampalayam, S., Moledina, N., Borland, A., Harding, S. A., Tsai, C.-J., & Leebens-Mack, J. (2019). Shared expression of crassulacean acid metabolism (CAM) genes pre-dates the origin of CAM in the genus Yucca. Journal of Experimental Botany, 70(22), 6597–6609.

Heyduk, K., Ray, J. N., & Leebens-Mack, J. (2021). Leaf anatomy is not correlated to CAM function in a C3+CAM hybrid species, Yucca gloriosa. Annals of Botany, 127(4), 437–449.

Hobo, T., Kowyama, Y., & Hattori, T. (1999). A bZIP factor, TRAB1, interacts with VP1 and mediates abscisic acid-induced transcription. Proceedings of the National Academy of Sciences of the United States of America, 96(26), 15348–15353.

Jung, C., Nguyen, N. H., & Cheong, J.-J. (2020). Transcriptional regulation of protein phosphatase 2C genes to modulate abscisic acid signaling. International Journal of Molecular Sciences, 21(24), 9517.

Katoh, K., Misawa, K., Kuma, K.-I., & Miyata, T. (2002). MAFFT: a novel method for rapid multiple sequence alignment based on fast Fourier transform. Nucleic Acids Research, 30(14), 3059–3066.

Konopka-Postupolska, D., Clark, G., Goch, G., Debski, J., Floras, K., Cantero, A., Fijolek, B., Roux, S., & Hennig, J. (2009). The role of annexin 1 in drought stress in Arabidopsis. Plant Physiology, 150(3), 1394–1410.

Kuznetsova, A., Brockhoff, P. B., & Christensen, R. H. B. (2017). lmerTest Package: Tests in Linear Mixed Effects Models. Journal of Statistical Software, 82, 1–26.

Láruson, Á. J., Yeaman, S., & Lotterhos, K. E. (2020). The Importance of Genetic Redundancy in Evolution. Trends in Ecology & Evolution, 35(9), 809–822.

Li, X., Ding, M., Wang, M., Yang, S., Ma, X., Hu, J., Song, F., Wang, L., & Liang, W. (2022). Proteome profiling reveals changes in energy metabolism, transport and antioxidation during drought stress in Nostoc flagelliforme. BMC Plant Biology, 22(1), 162.

Liao, Y., Smyth, G. K., & Shi, W. (2014). FeatureCounts: An efficient general purpose program for assigning sequence reads to genomic features. Bioinformatics, 30(7), 923–930. arXiv.

Loewus, F. A., & Murthy, P. P. N. (2000). myo-Inositol metabolism in plants. Plant Science: An International Journal of Experimental Plant Biology, 150(1), 1–19.

Lüttge, U. (1987). Carbon dioxide and water demand: Crassulacean acid metabolism (cam), a versatile ecological adaptation exemplifying the need for integration in ecophysiological work. The New Phytologist, 106(4), 593–629.

Magwanga, R. O., Lu, P., Kirungu, J. N., Lu, H., Wang, X., Cai, X., Zhou, Z., Zhang, Z., Salih, H., Wang, K., & Liu, F. (2018). Characterization of the late embryogenesis abundant (LEA) proteins family and their role in drought stress tolerance in upland cotton. BMC Genetics, 19(1), 6.

Males, J. (2018). Concerted anatomical change associated with crassulacean acid metabolism in the Bromeliaceae. Functional Plant Biology: FPB, 45(7), 681–695.

Martin, C. E. (1994). Physiological ecology of the Bromeliaceae. The Botanical Review; Interpreting Botanical Progress, 60(1), 1–82.

Martin, C. E., Hsu, R. C.-C., & Lin, T.-C. (2009). The relationship between CAM and leaf succulence in two epiphytic vines, Hoya carnosa and Dischidia formosana (Asclepiadaceae), in a subtropical rainforest in northeastern Taiwan. Photosynthetica, 47(3), 445–450.

Mendoza, C. G., Granados-Aguilar, X., Donadío, S., Salazar, G. A., Flores-Cruz, M., Hágsater, E., Starr, J. R., Ibarra-Manríquez, G., Fragoso-Martínez, I., & Magallón, S. (2017). Geographic structure in two highly diverse lineages of Tillandsia (Bromeliaceae). Botany, 95(7), 641–651.

Ming, R., VanBuren, R., Wai, C. M., Tang, H., Schatz, M. C., Bowers, J. E., Lyons, E., Wang, M. L., Chen, J., Biggers, E., Zhang, J., Huang, L., Zhang, L., Miao, W., Zhang, J., Ye, Z., Miao, C., Lin, Z., Wang, H., … Yu, Q. (2015). The pineapple genome and the evolution of CAM photosynthesis. Nature Genetics, 47(12), 1435–1442.

Minh, B. Q., Schmidt, H. A., Chernomor, O., Schrempf, D., Woodhams, M. D., von Haeseler, A., & Lanfear, R. (2020). IQ-TREE 2: New Models and Efficient Methods for Phylogenetic Inference in the Genomic Era. Molecular Biology and Evolution, 37(5), 1530–1534.

Mioto, P. T., & Mercier, H. (2013). Abscisic acid and nitric oxide signaling in two different portions of detached leaves of Guzmania monostachia with CAM up-regulated by drought. Journal of Plant Physiology, 170(11), 996–1002.

Müller, M., Seifert, S., Lübbe, T., Leuschner, C., & Finkeldey, R. (2017). De novo transcriptome assembly and analysis of differential gene expression in response to drought in European beech. PloS One, 12(9), e0184167.

Neales, T. F. (1975). The gas exchange patterns of CAM plants. In Environmental and Biological Control of Photosynthesis (pp. 299–310). Springer Netherlands.

Nelson, E. A., & Sage, R. F. (2008). Functional constraints of CAM leaf anatomy: tight cell packing is associated with increased CAM function across a gradient of CAM expression. Journal of Experimental Botany, 59(7), 1841–1850.

Nelson, E. A., Sage, T. L., & Sage, R. F. (2005). Functional leaf anatomy of plants with crassulacean acid metabolism. Functional Plant Biology: FPB, 32(5), 409–419.

Nishizawa, A., Yabuta, Y., & Shigeoka, S. (2008). Galactinol and raffinose constitute a novel function to protect plants from oxidative damage. Plant Physiology, 147(3), 1251–1263.

Nowak, E. J., & Martin, C. E. (1997). Physiological and Anatomical Responses to Water Deficits in the Cam Epiphyte Tillandsia ionantha (Bromeliaceae). International Journal of Plant Sciences, 158(6), 818–826.

Nowak, M. A., Boerlijst, M. C., Cooke, J., & Smith, J. M. (1997). Evolution of genetic redundancy. Nature, 388(6638), 167–171.

Ogburn, R. M., & Edwards, E. J. (2009). Anatomical variation in Cactaceae and relatives: Trait lability and evolutionary innovation. American Journal of Botany, 96(2), 391–408.

Ohrui, T., Nobira, H., Sakata, Y., Taji, T., Yamamoto, C., Nishida, K., Yamakawa, T., Sasuga, Y., Yaguchi, Y., Takenaga, H., & Tanaka, S. (2007). Foliar trichome- and aquaporin-aided water uptake in a drought-resistant epiphyte Tillandsia ionantha Planchon. Planta, 227(1), 47–56.

Osmond, C. B. (1978). Crassulacean Acid Metabolism: A Curiosity in Context. Annual Review of Plant Physiology, 29(1), 379–414.

Pierce, S., Winter, K., & Griffiths, H. (2002). Carbon isotope ratio and the extent of daily CAM use by Bromeliaceae. The New Phytologist, 156(1), 75–83.

Quezada, I. M., & Gianoli, E. (2011). Crassulacean acid metabolism photosynthesis in Bromeliaceae: an evolutionary key innovation. Biological Journal of the Linnean Society. Linnean Society of London, 104(2), 480–486.

Robinson, M. D., McCarthy, D. J., & Smyth, G. K. (2009). edgeR: A Bioconductor package for differential expression analysis of digital gene expression data. Bioinformatics, 26(1), 139–140.

Robinson, M. D., & Oshlack, A. (2010). A scaling normalization method for differential expression analysis of RNA-seq data. Genome Biology, 11(3), R25.

Schiller, K., & Bräutigam, A. (2021). Engineering of Crassulacean Acid Metabolism. Annual Review of Plant Biology, 72, 77–103.

Schubert, M., Lindgreen, S., & Orlando, L. (2016). AdapterRemoval v2: rapid adapter trimming, identification, and read merging. BMC Research Notes, 9(1), 88.

Silvera, K., Santiago, L. S., Cushman, J. C., & Winter, K. (2009). Crassulacean acid metabolism and epiphytism linked to adaptive radiations in the Orchidaceae. Plant Physiology, 149(4), 1838–1847.

Silvera, K., Santiago, L. S., & Winter, K. (2005). Distribution of crassulacean acid metabolism in orchids of Panama: evidence of selection for weak and strong modes. Functional Plant Biology: FPB, 32(5), 397–407.

Steiner, C. C., Römpler, H., Boettger, L. M., Schöneberg, T., & Hoekstra, H. E. (2009). The genetic basis of phenotypic convergence in beach mice: similar pigment patterns but different genes. Molecular Biology and Evolution, 26(1), 35–45.

Stiles, K. C., & Martin, C. E. (1996). Effects of drought stress on CO2 exchange and water relations in the CAM epiphyte Tillandsia utriculata (Bromeliaceae). Journal of Plant Physiology, 149(6), 721–728.

Szukala, A., Lovegrove-Walsh, J., Luqman, H., Fior, S., Wolfe, T. M., Frajman, B., Schönswetter, P., & Paun, O. (2023). Polygenic routes lead to parallel altitudinal adaptation in Heliosperma pusillum (Caryophyllaceae). Molecular Ecology, 32(8), 1832–1847.

Ting, I. P. (1981). Effects of abscisic acid on CAM in Portulacaria afra. Photosynthesis Research, 2(1), 39–48.

Vahisalu, T., Kollist, H., Wang, Y.-F., Nishimura, N., Chan, W.-Y., Valerio, G., Lamminmäki, A., Brosché, M., Moldau, H., Desikan, R., Schroeder, J. I., & Kangasjärvi, J. (2008). SLAC1 is required for plant guard cell S-type anion channel function in stomatal signalling. Nature, 452(7186), 487–491.

Vera-Paz, S. I., Granados Mendoza, C., Díaz Contreras Díaz, D. D., Jost, M., Salazar, G. A., Rossado, A. J., Montes-Azcué, C. A., Hernández-Gutiérrez, R., Magallón, S., Sánchez-González, L. A., Gouda, E. J., Cabrera, L. I., Ramírez-Morillo, I. M., Flores-Cruz, M., Granados-Aguilar, X., Martínez-García, A. L., Hornung-Leoni, C. T., Barfuss, M. H. J., & Wanke, S. (2023). Plastome phylogenomics reveals an early Pliocene North- and Central America colonization by long-distance dispersal from South America of a highly diverse bromeliad lineage. Frontiers in Plant Science, 14, 1205511.

Wai, C. M., Weise, S. E., Ozersky, P., Mockler, T. C., Michael, T. P., & VanBuren, R. (2019). Time of day and network reprogramming during drought induced CAM photosynthesis in Sedum album. PLoS Genetics, 15(6), e1008209.

Wan, X., Mo, A., Liu, S., Yang, L., & Li, L. (2011). Constitutive expression of a peanut ubiquitin-conjugating enzyme gene in Arabidopsis confers improved water-stress tolerance through regulation of stress-responsive gene expression. Journal of Bioscience and Bioengineering, 111(4), 478–484.

Wasternack, C. (2014). Action of jasmonates in plant stress responses and development--applied aspects. Biotechnology Advances, 32(1), 31–39.

Wickell, D., Kuo, L.-Y., Yang, H.-P., Dhabalia Ashok, A., Irisarri, I., Dadras, A., de Vries, S., de Vries, J., Huang, Y.-M., Li, Z., Barker, M. S., Hartwick, N. T., Michael, T. P., & Li, F.-W. (2021). Underwater CAM photosynthesis elucidated by Isoetes genome. Nature Communications 2021 12:1, 12(1), 1–13.

Wilkinson, S., & Davies, W. J. (2002). ABA-based chemical signalling: the co-ordination of responses to stress in plants. *Plant*, Cell & Environment, 25(2), 195–210.

Winter, K. (2019). Ecophysiology of constitutive and facultative CAM photosynthesis. Journal of Experimental Botany. 10.1093/jxb/erz002

Winter, K., & Holtum, J. A. M. (2002). How closely do the delta(13)C values of Crassulacean Acid metabolism plants reflect the proportion of CO(2) fixed during day and night? Plant Physiology, 129(4), 1843–1851.

Winter, K., & Holtum, J. A. M. (2014). Facultative crassulacean acid metabolism (CAM) plants: powerful tools for unravelling the functional elements of CAM photosynthesis. Journal of Experimental Botany, 65(13), 3425–3441.

Winter, K., & Smith, J. A. C. (2022). CAM photosynthesis: the acid test. The New Phytologist, 233(2), 599–609.

Winter, K., & Ziegler, H. (1992). Induction of crassulacean acid metabolism in Mesembryanthemum crystallinum increases reproductive success under conditions of drought and salinity stress. Oecologia, 92(4), 475–479.

Yardeni, G., Barfuss, M. H. J., Till, W., Thornton, M. R., Crego, C. G., Lexer, C., Leroy, T., & Paun, O. (2025). The explosive radiation of the Neotropical Tillandsia subgenus Tillandsia (Bromeliaceae) has been accompanied by pervasive hybridization. *Systematic Biology*, syaf039.

Yardeni, G., Viruel, J., Paris, M., Hess, J., Groot Crego, C., de La Harpe, M., Rivera, N., Barfuss, M. H. J., Till, W., Guzmán-Jacob, V., Krömer, T., Lexer, C., Paun, O., & Leroy, T. (2021). Taxon-specific or universal? Using target capture to study the evolutionary history of rapid radiations. Molecular Ecology Resources. 10.1111/1755-0998.13523

Yeaman, S., Hodgins, K. A., Lotterhos, K. E., Suren, H., Nadeau, S., Degner, J. C., Nurkowski, K. A., Smets, P., Wang, T., Gray, L. K., Liepe, K. J., Hamann, A., Holliday, J. A., Whitlock, M. C., Rieseberg, L. H., & Aitken, S. N. (2016). Convergent local adaptation to climate in distantly related conifers. Science, 353(6306), 1431–1433.

Zhu, J.-K. (2002). Salt and drought stress signal transduction in plants. Annual Review of Plant Biology, 53, 247–273.

