## Supplementary figures for "Divergent transcriptional architectures beyond core CAM genes in facultative and constitutive CAM species in Tillandsia"

### Divergent transcriptional architecture beyond core CAM genes in closely related facultative and constitutive CAM species of the subgenus Tillandsia

C. Groot Crego, S. Saadain, M.de La Harpe, J. Hess, M.H.J. Barfuss, W. Till, C. Lexer, O. Paun

Figure S1: Diagnostic plots for Linear Mixed-Effects Models constructed in each species to assess the effect of time and watering regime on titratable acidity measurements. Column-wise, the plots are organised per species, as a separate model was constructed for each. The first row of plots displays the scatter of residuals to the fitted values, while the second row shows the QQ-plot and the results of the Shapiro-Wilk test on the residuals.

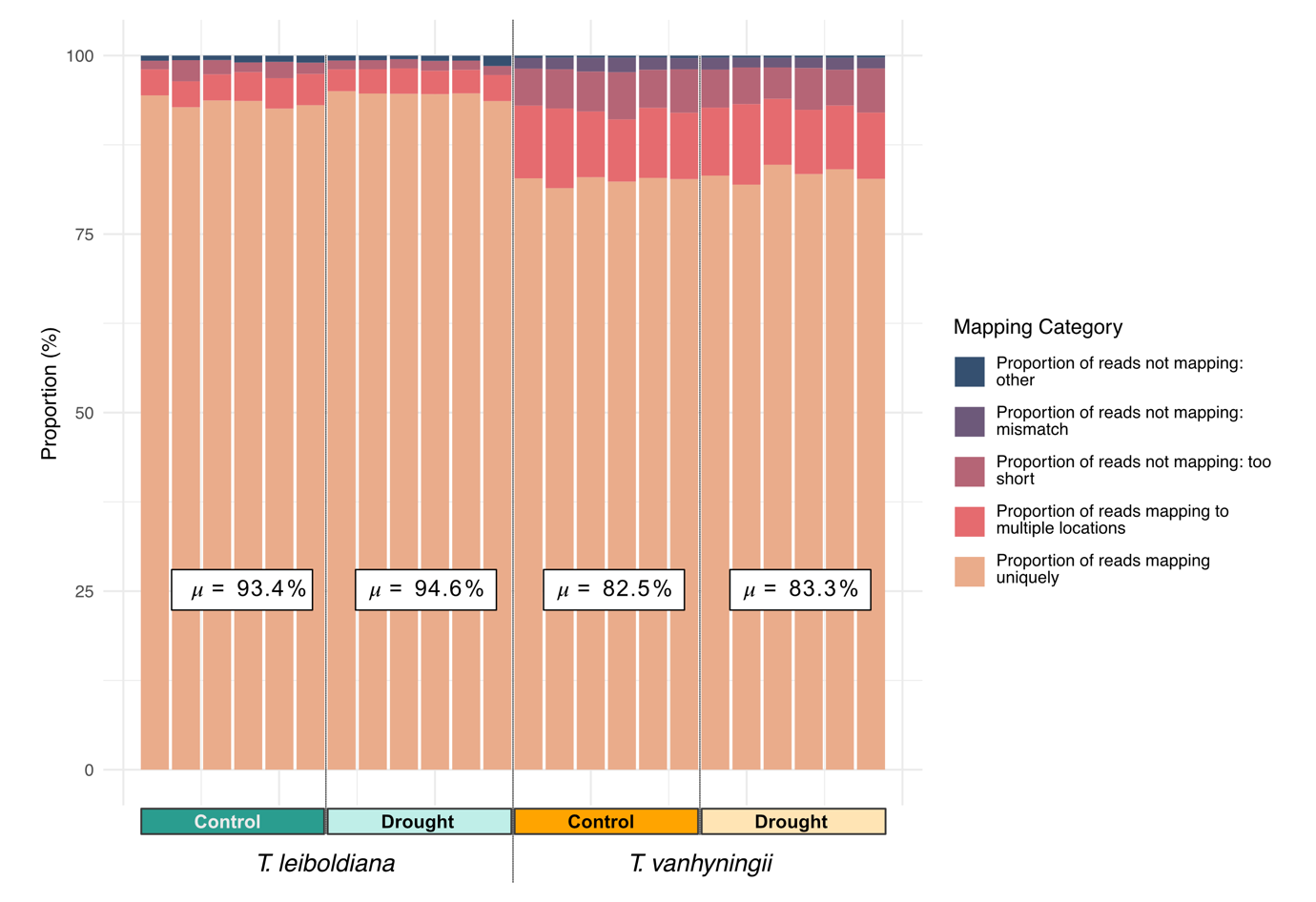

Figure S2: Mapping rates across 24 samples of different species and watering regimes. The stacked bar plot specifies the proportion of reads that mapped uniquely, to multiple locations or that were unmapped for several reasons. Average unique mapping rates are given for each group of samples of a given species and watering regime. T. leiboldiana reads were mapped to the conspecific T. leiboldiana reference genome, while T. vanhyningii reads were mapped to the reference genome of its close relative, T. fasciculata.

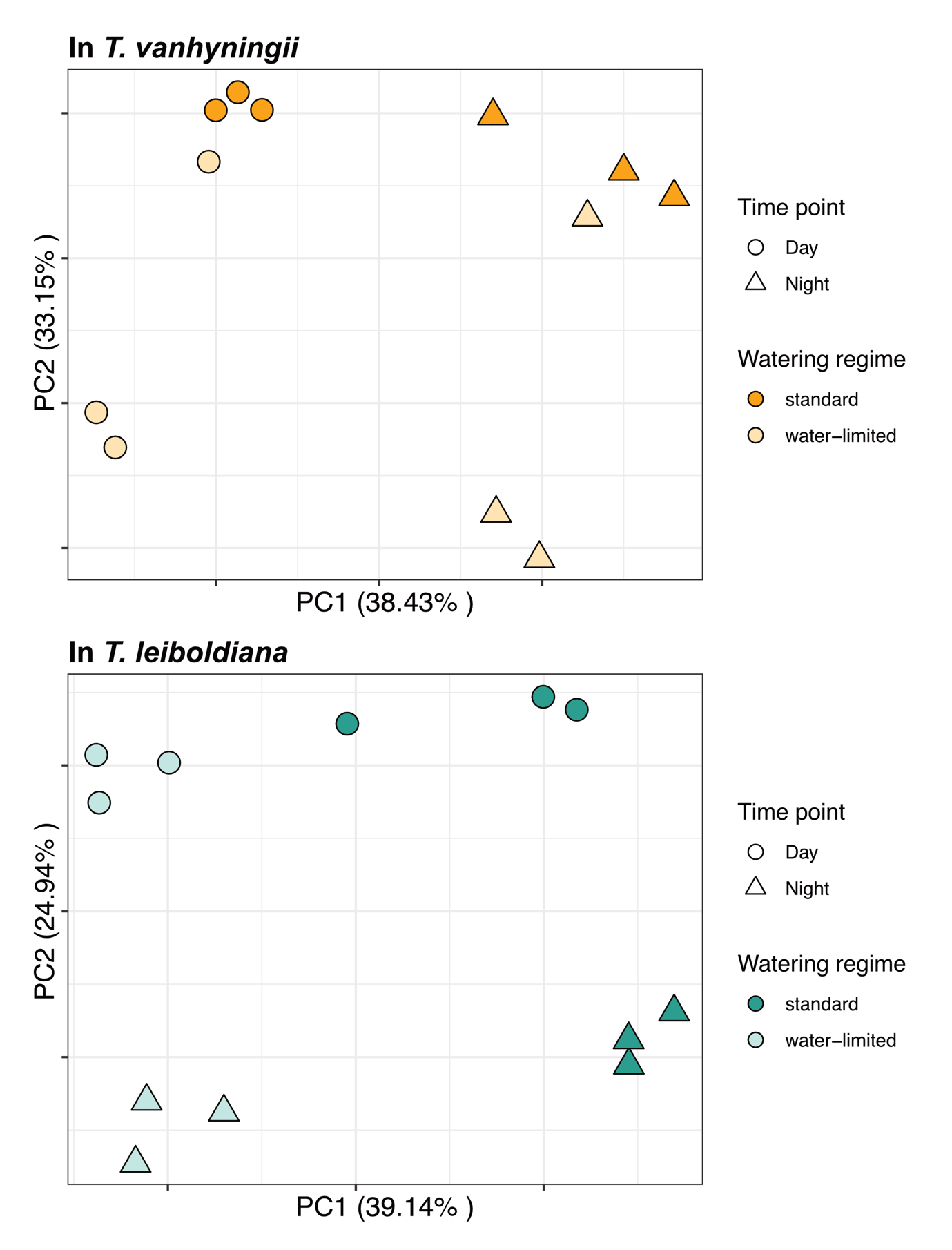

Figure S3: Principal component analysis of TMM-normalised read counts in log(CPM).

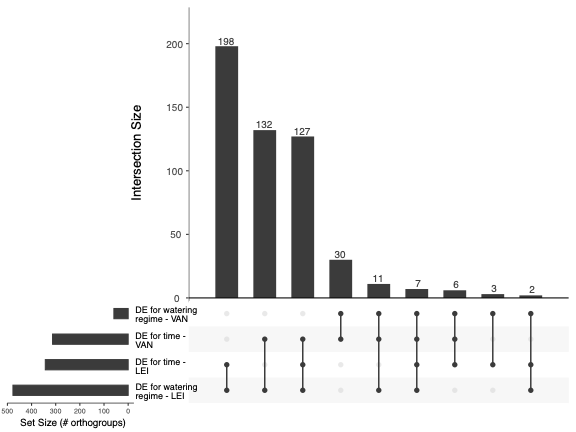
Figure S4: Overview of the overlap between DE analyses across time points and across watering regimes. The bar plots represent the number of orthogroups overlapping between DE analyses, sorted in descending order. Below, the linked dots represent the analyses which display each overlap. The vertically oriented bar plot on the left-side shows the total number of DE orthogroups called in each analysis. DE analyses were performed between time points and watering regimes in each species (VAN = T. vanhyningii, LEI = T. leiboldiana)

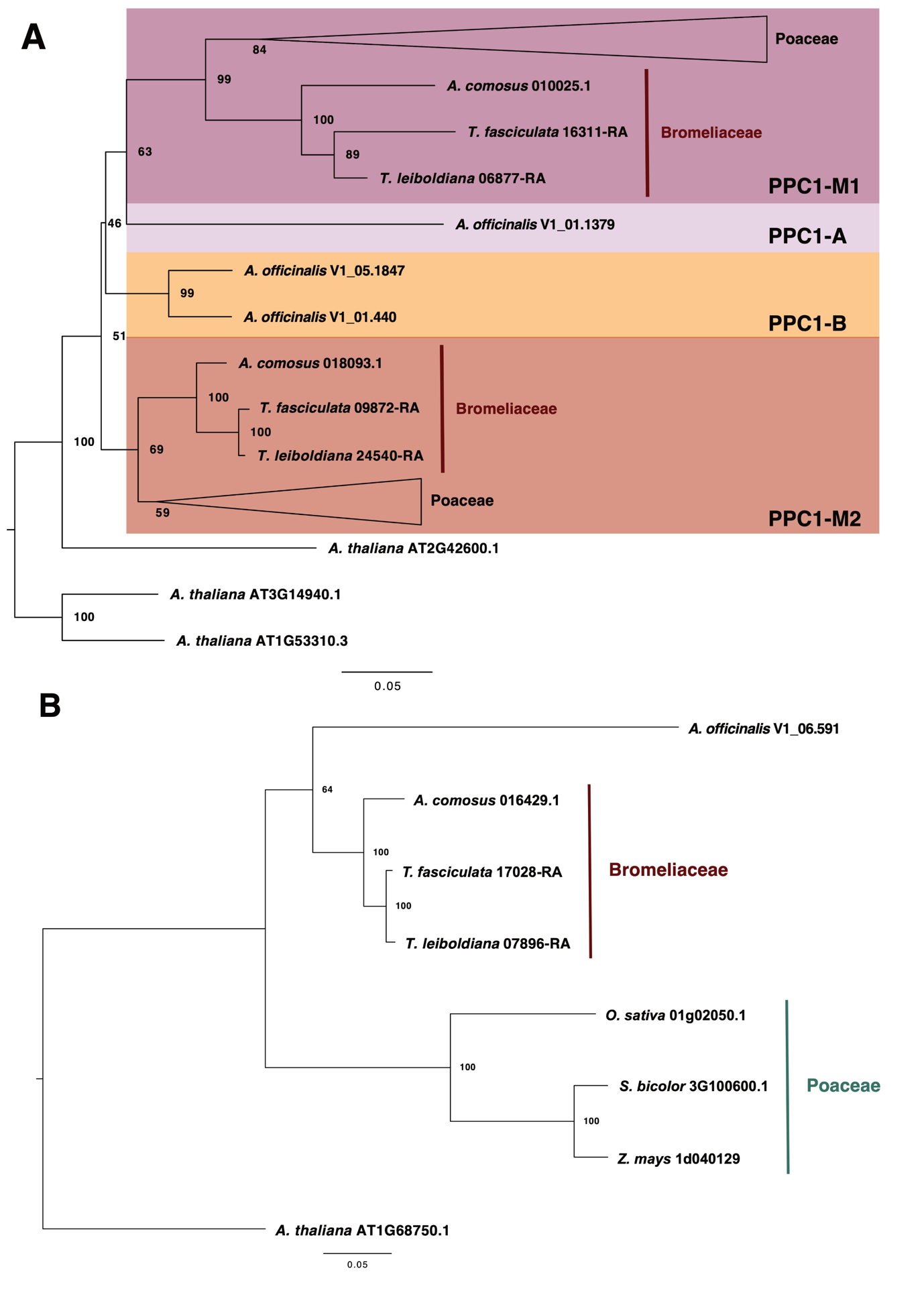

Figure S5: Gene trees of PPC1 (A) and PPC2 (B).

Figure S6: Gene trees of PPCK with the isozymes PPCK1 and PPCK2.

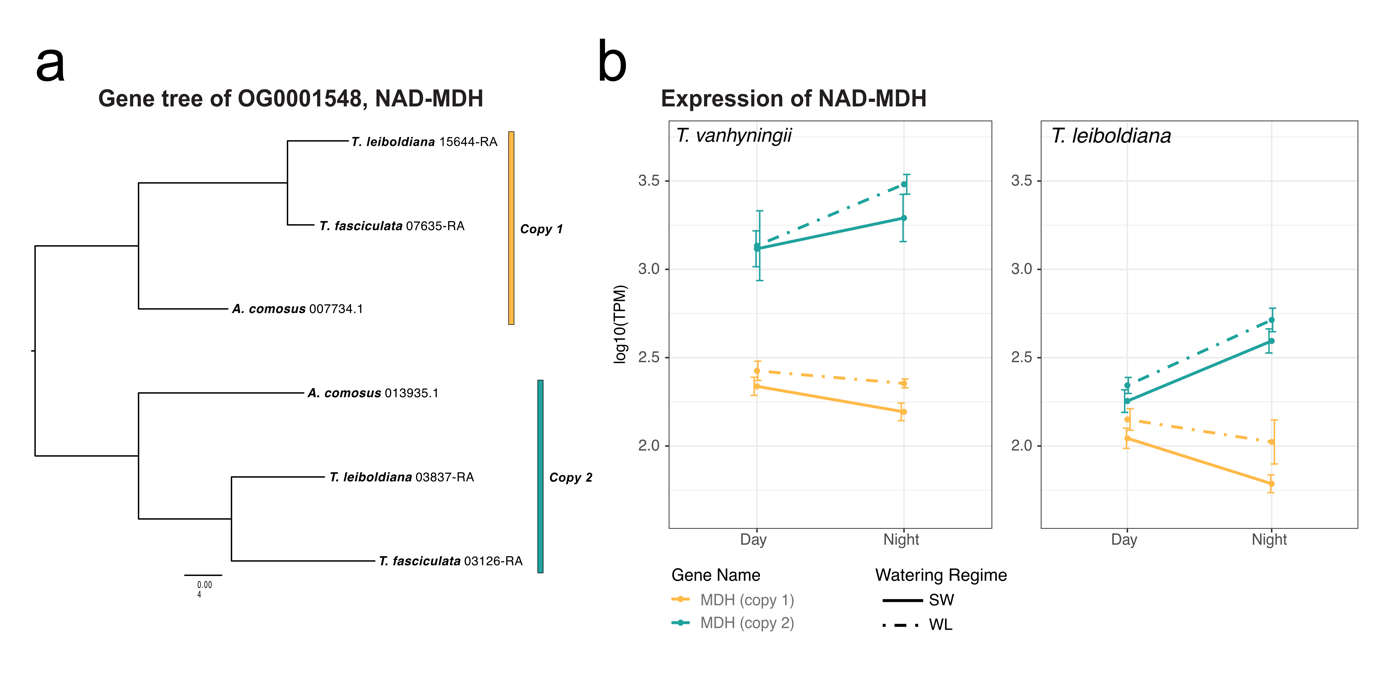

Figure S7: Expression of MDH orthologs. A) Gene tree of orthogroup OG0001548, which encodes for an NAD-dependent malate dehydrogenase, obtained from Groot Crego et al. 2024. B) Transcript abundance in “Transcripts per Million” (TPM) for all gene copies belonging to OG0001548. TPM values are transformed to a logarithmic scale.

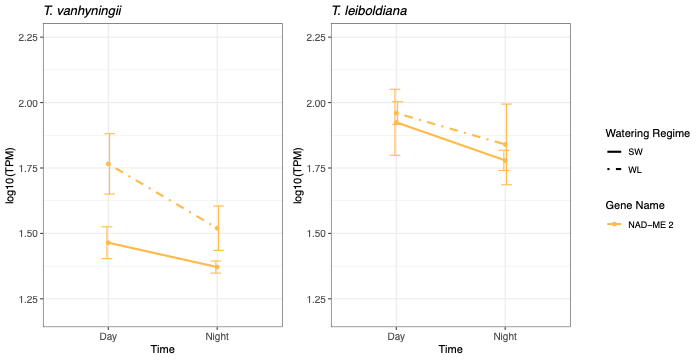

Figure S8: Expression of malic enzyme (NAD-ME 2). Transcript abundance in “Transcripts per Million” (TPM) for NAD-ME in T. vanhyningii and T. leiboldiana at two time points under standard and water-limited conditions. TPM values are transformed to a logarithmic scale.

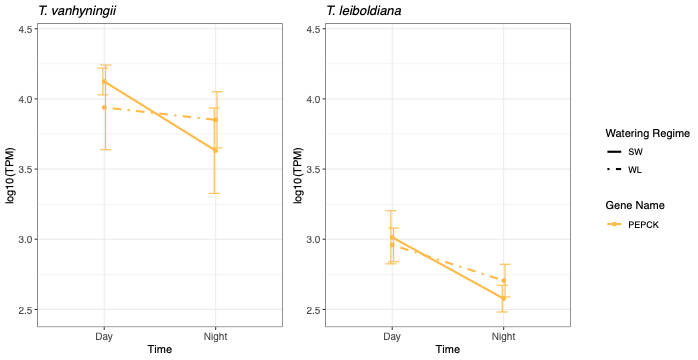

Figure S9: Expression of Phosphoenolpyruvate carboxykinase (PEPCK). Transcript abundance in “Transcripts per Million” (TPM) for NAD-ME in T. vanhyningii and T. leiboldiana at two time points under standard and water-limited conditions. TPM values are transformed to a logarithmic scale.

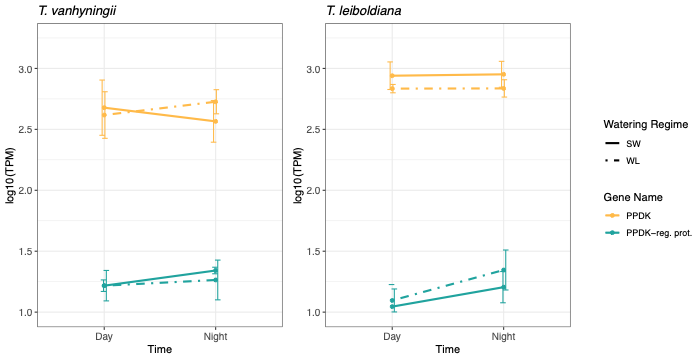

Figure S10: Expression of pyruvate-phosphate dikinase (PPDK) and PPDK regulatory protein. Transcript abundance in “Transcripts per Million” (TPM) for NAD-ME in T. vanhyningii and T. leiboldiana at two time points under standard and water-limited conditions. TPM values are transformed to a logarithmic scale.

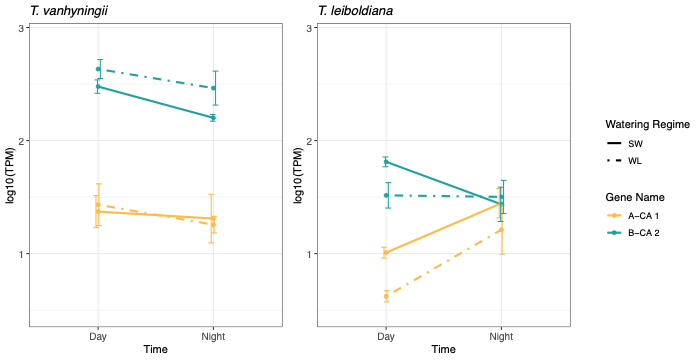

Figure S11: Expression of carbonic anhydrases. Transcript abundance in “Transcripts per Million” (TPM) for α-CA and β-CA in T. vanhyningii and T. leiboldiana at two time points under standard and water-limited conditions. TPM values are transformed to a logarithmic scale. α-CA was DE in T. leiboldiana under water-limited conditions.

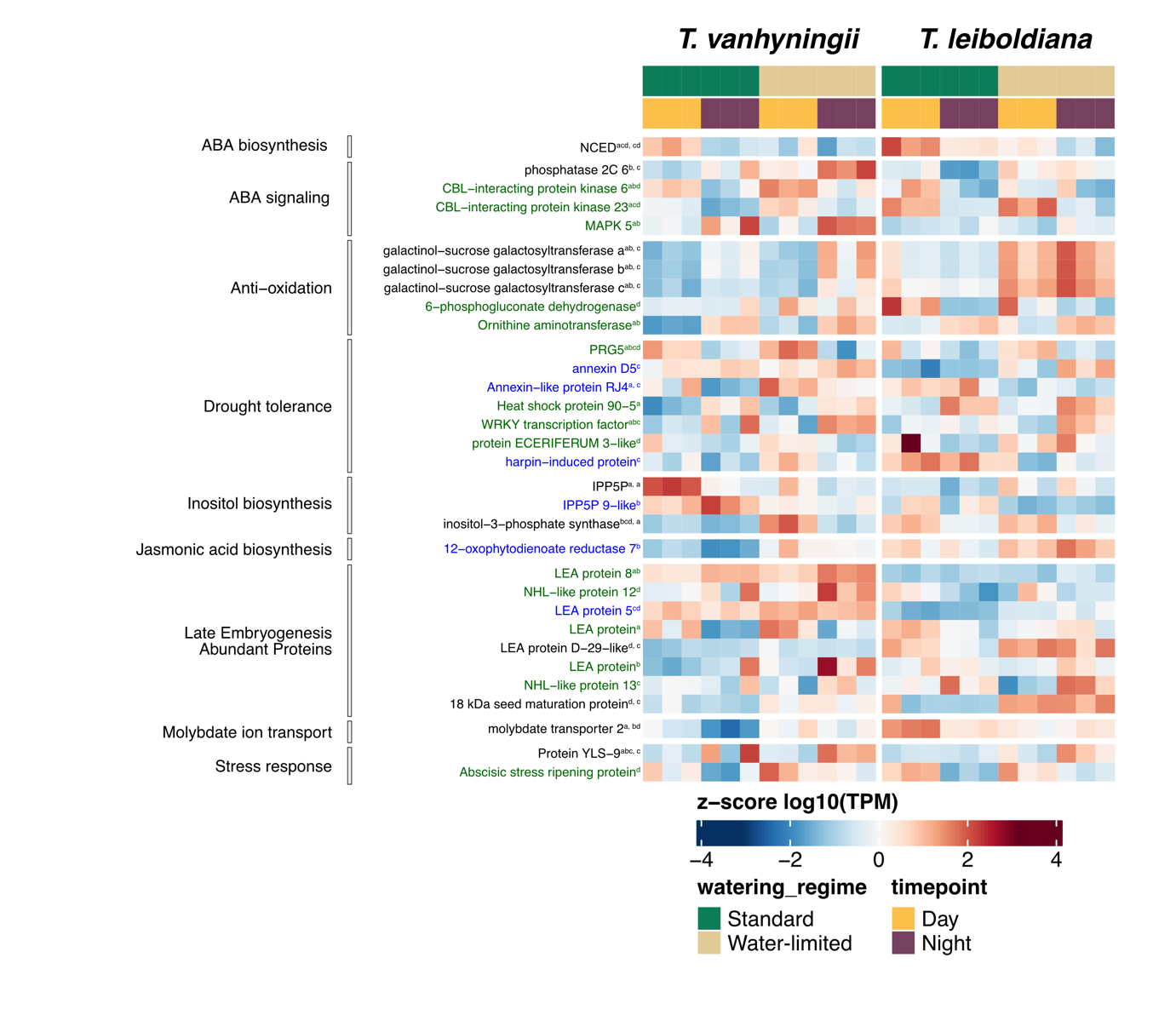
Figure S12: Transcript abundance in “Transcripts per Million” (TPM) for a subset of orthogroups related to drought response in T. vanhyningii (left panel, using the T. fasciculata ortholog) and T. leiboldiana (right panel, using the T. leiboldiana ortholog) at two timepoints and watering regimes. TPM values are transformed to a logarithmic scale and centered as a z-score. Displayed genes are organised in functional categories and gene name colour reflects their differential expression status: blue = DE between watering regimes, green = DE between timepoints, black is DE in both comparisons and grey = not DE. The letters in superscript indicate the intersection of the Venn diagram that each orthogroup belongs to. When an orthogroup is DE in both comparisons, the first superscripted letter indicates the intersection of the timewise Venn diagram (Figure 4a), and the second superscripted letter, separated by a comma, indicates the intersection of the Venn diagram for watering regimes (Figure 4b). Abbreviations: PRG - Protein PROTON GRADIENT REGULATION; NHL - NDR1/HIN1-like; MAPK - mitogen-activated protein kinase; IPP5P = Inositol Polyphosphate 5-Phosphatase

**Supplementary Note: Comparison of results using a conspecific versus a non-conspecific reference genome**

Here we provide a comparison of DGE results stemming from RNA-seq mapping of T. leiboldiana reads to a conspecific reference and a non-conspecific genome (T. fasciculata).

1. **Number of Up- and down-regulated genes:**

When comparing the numbers of DE genes that are up- and downregulated, as shown in the table below, the difference between approaches is small. At most, 30 more DE genes between watering regimes are discovered by mapping T. leiboldiana conspecifically, yet this is only a 5 % difference.

|  | Number of upregulated genes | | | Number of downregulated genes | | |
| --- | --- | --- | --- | --- | --- | --- |
|  | Mapped to Tfas | Mapped to Tlei | Difference | Mapped to Tfas | Mapped to Tlei | Difference |
| Day vs Night in SW | 615 | 603 | **-12** | 254 | 263 | **9** |
| Day vs Night in WL | 447 | 428 | **-19** | 392 | 370 | **-22** |
| WL vs SW during the Day | 75 | 73 | **-2** | 326 | 313 | **-13** |
| WL vs SW during the Night | 557 | 587 | **30** | 532 | 540 | **8** |

2. **Number and overlap of orthogroups**

To see how similar the composition of DE genes is between the two approaches, we looked at the overlap of orthogroup occurrence in each DE comparison. Across all comparisons, between 80-90 % of orthogroups with DE genes were called with both approaches. About 10-20 % of orthogroups are therefore only recovered as having a DE gene in one mapping approach. This is in part due to an added layer of complexity stemming from the orthology analysis (not all orthologous / paralogous relations were resolved), but also due to copy number differences between species. However, this is only affecting a minority of orthogroups.

|  | Number of orthogroups | | |
| --- | --- | --- | --- |
|  | Mapped to Tfas | Mapped to Tlei | Overlap |
| Day vs Night in SW | 763 | 737 | **627** |
| Day vs Night in WL | 716 | 716 | **619** |
| WL vs SW during the Day | 359 | 346 | **281** |
| WL vs SW during the Night | 941 | 1004 | **787** |

3. **Differences in Venn diagrams on the genic and orthogroup level between species**

When comparing the Venn diagrams resulting from the two mapping approaches, very little changes in terms of the relative composition of the Venn diagrams, and therefore also regarding our conclusions based on overlap between species.

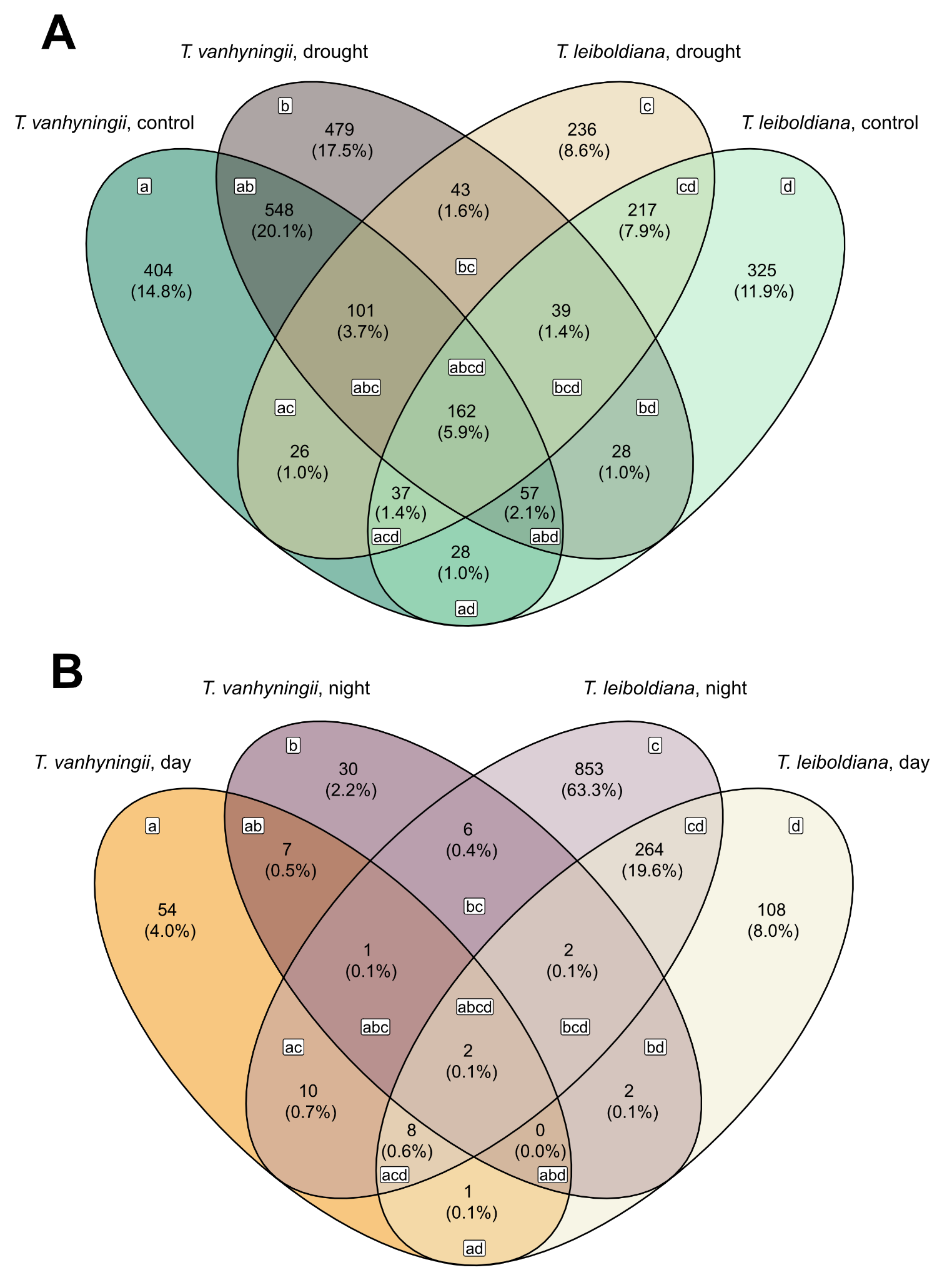

Venn diagrams showing overlap of DE genes when mapping both species to the T. fasciculata reference genome

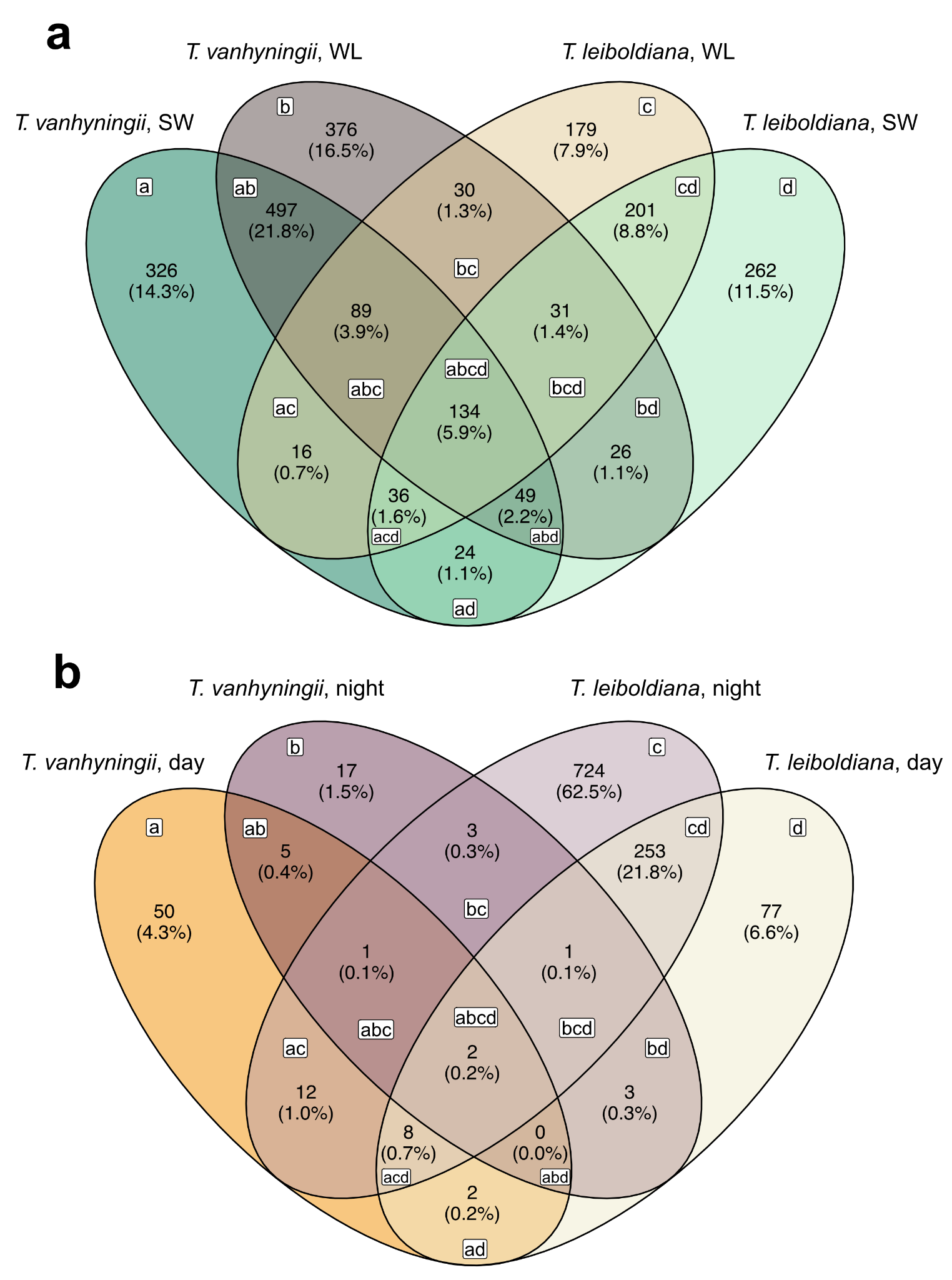

Venn diagrams showing overlap of DE genes on the orthogroup level when mapping T. leiboldiana to its conspecific reference genome

4. **Presence and overlap of CAM-related genes**

The number of CAM-related genes in each intersection of the Venn-diagrams is largely similar between approaches.

| Day vs Night | | | |
| --- | --- | --- | --- |
| Nr. of CAM genes | | Venn diagram intersection | Group description |
| Using a non-conspecific genome | Using a conspecific genome for T. leiboldiana |  |  |
| 27 | 27 | a | Unique VAN control |
| 21 | 20 | b | Unique VAN drought |
| 8 | 9 | c | Unique LEI drought |
| 12 | 13 | d | Unique LEI control |
| 5 | 6 | ab | VAN drought + control |
| 2 | 2 | ac | VAN control + LEI drought |
| 5 | 5 | ad | VAN + LEI control |
| 2 | 3 | bc | VAN + LEI drought |
| 15 | 17 | bd | VAN drought + LEI control |
| 1 | 3 | cd | LEI drought + control |
| 1 | 2 | abc | VAN + LEI drought |
| 2 | 2 | abd | VAN + LEI control |
| 17 | 15 | acd | VAN control + LEI |
| 13 | 13 | bcd | VAN drought + LEI |
| 18 | 15 | abcd | all groups |
| Standard watering vs water-limited conditions | | | |
| Nr. of CAM genes | | Venn diagram intersection | Group description |
| Using a non-conspecific genome | Using a conspecific genome for T. leiboldiana |  |  |
| 2 | 5 | a | Unique VAN day |
| 1 | 1 | b | Unique VAN night |
| 0 | 1 | c | Unique LEI night |
| 0 | 1 | d | Unique LEI day |
| 0 | 2 | ab | VAN day + night |
| 0 | 1 | ac | VAN day + LEI night |
| 1 | 2 | ad | VAN day + LEI day |
| 0 | 2 | bc | VAN night + LEI night |
| 3 | 1 | bd | VAN night + LEI day |
| 0 | 1 | cd | LEI day + LEI night |
| 0 | 1 | abc | VAN + LEI night |
| 0 | 1 | abd | VAN + LEI day |
| 51 | 57 | acd | VAN day + LEI |
| 21 | 21 | bcd | VAN night + LEI |
| 5 | 2 | abcd | all groups |

5. **Expression of PPCK and PPC**

The expression levels of both gene families remain largely unchanged between mapping approaches. This is also the case for CAM-related genes shown in Fig. S4 (comparison not shown here).

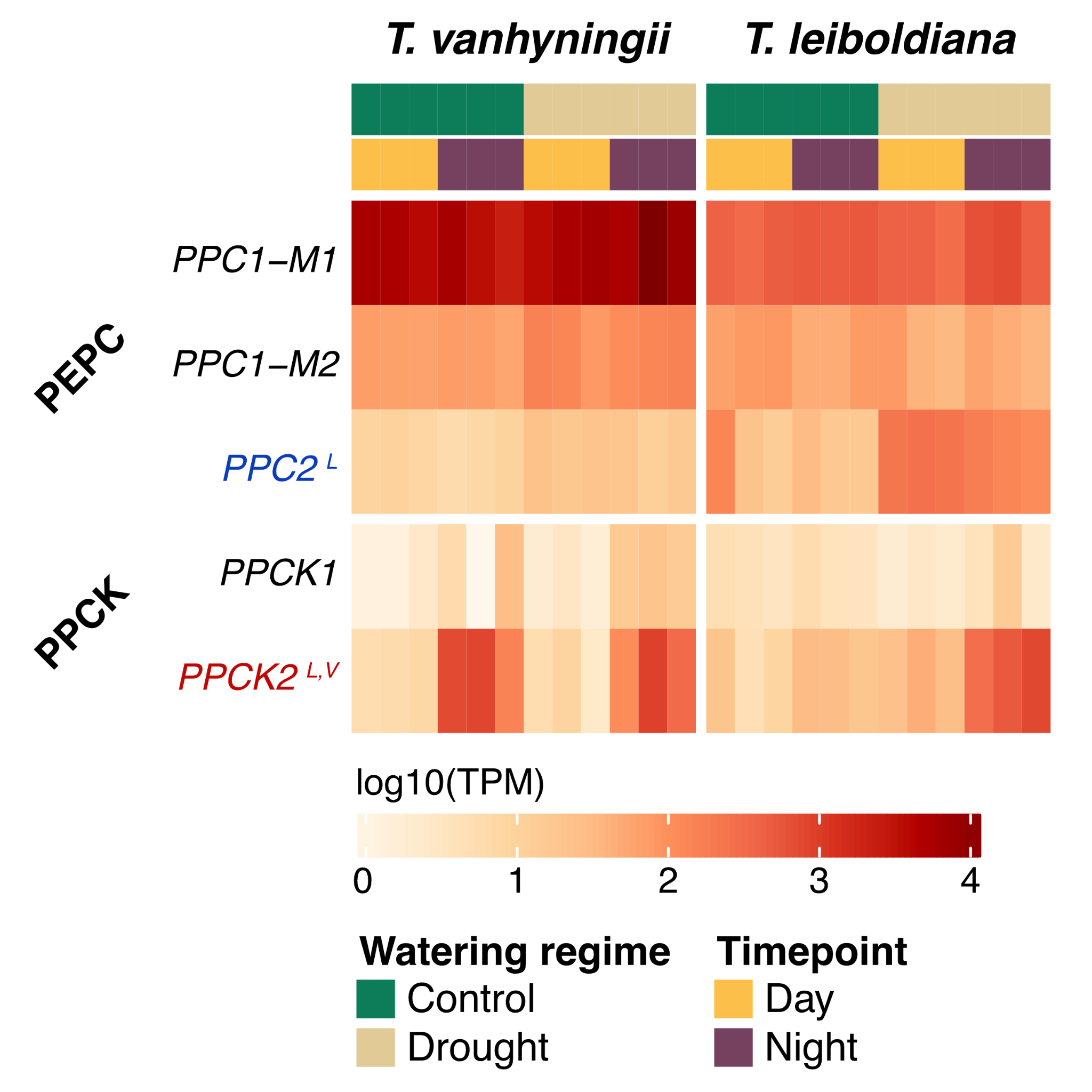

Expression of PEPC and PPCK genes when using T. fasciculata as reference genome

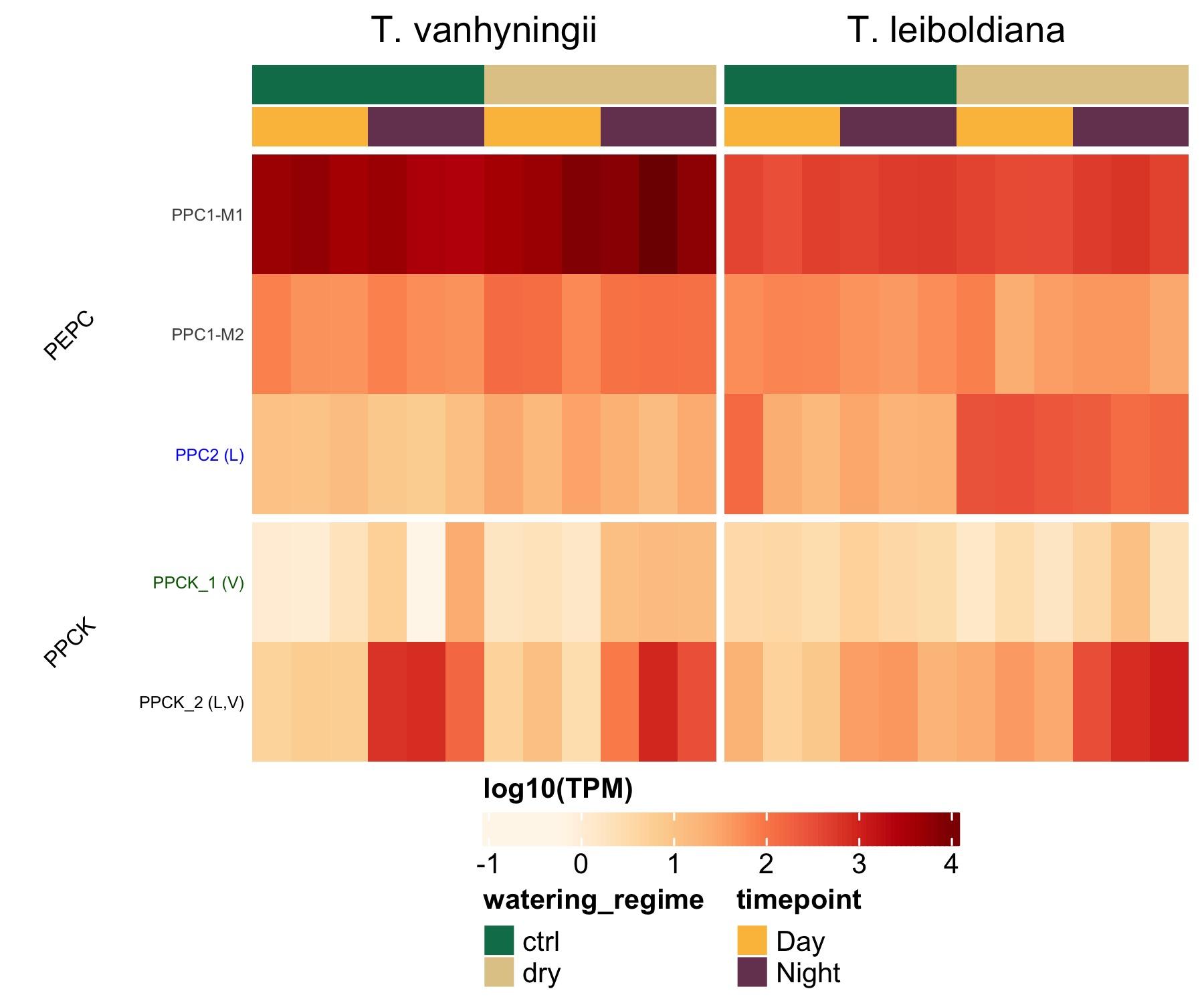

Expression of PEPC and PPCK genes when using T. leiboldiana as reference genome for T. leiboldiana and T. fasciculata for T. vanhyningii.

These comparisons show that while using a non-conspecific reference genome for these analyses will slightly change which genes are recovered as DE, the overall results presented in our paper change very little between approaches. Additionally, T. vanhyningii is more closely related to T. fasciculata than T. leiboldiana, as stated in the methods section. Therefore, any differences in DE expression results in T. vanhyningii stemming from using T. fasciculata as reference genome, rather than a conspecific genome, are expected to be fewer and smaller than the differences we are witnessing in the comparisons with T. leiboldiana.
